# A Multi-Model Ensemble Reveals Soil Carbon Gains from Regenerative Practices in the U.S. Midwest Cropland

**DOI:** 10.1101/2025.02.04.636509

**Authors:** Bruno Basso, Tommaso Tadiello, Neville Millar, G. Philip Robertson, Keith Paustian, Fidel S. Maureira, Susana Albarenque, Brian Baer, Lydia Price, Prateek Sharma, Chris Villalobos, Ames Fowler, Mathieu Delandmeter, Marco Acutis, Sotirios Archontoulis, Kristofer R. Covey, Luca Doro, Benjamin Dumont, Peter R. Grace, Gerrit Hoogenboom, James W. Jones, Alessia Perego, Alexander Ruane, Claudio O. Stöckle, Yao Zhang

## Abstract

Process-based cropping systems models (CSMs) are key components of measurement, monitoring, reporting, and verification (MMRV) frameworks of carbon markets, but their application suffers from model-specific differences that keep any one model from working well across all combinations of soils, climates, crops, and agronomic practices at varying scales. Multi-model ensemble (MME), successfully used to quantify soil, management and climate impact on crop productivity, provide an opportunity to better estimate changes in soil organic carbon (SOC) outcomes for agronomic practices that have the potential to mitigate SOC loss at scale. We used an MME across 46 million hectares of US Midwest cropland at a resolution of 4- km^2^ to assess the aggregate ability of different regenerative practices to sequester SOC at this scale compared to their dynamic baselines. MME was validated with long-term experimental data and compared to its constituent CSMs, showing greater accuracy and lower uncertainty. The results show that adopting no-till combined with cover crops increased SOC stocks by 0.36 ± 0.12 Mg ha^-1^ yr^-1^ aggregated across the entire U.S. Midwest cropland. At the regional scale, this corresponds to a net SOC gain of 16.4 Tg C yr^-1^ compared to business-as-usual baselines. These benefits are approximately halved when each management change is practiced individually, and the modest gains are only fully realized when continued over the long-term in soils with low initial carbon stock. Results demonstrate the power of MMEs run at high resolution for providing robust estimates of environmental outcomes following agricultural practice change, and for pinpointing locations for most effective intervention. This approach can alleviate many producer carbon market participation barriers and help address market issues while ultimately supporting large-scale regenerative agriculture initiatives.

## 1. Introduction

Soil organic carbon (SOC) storage is a key strategy for agricultural climate change mitigation but implementing reliable SOC-building practices on croplands across the landscape faces considerable challenges: inherent field variability, numerous quantification approaches, practical and financial barriers, and vested interests limit data accuracy, reproducibility, and scalability. Uncoordinated, inconsistent carbon market initiatives reflect these challenges (Wongpiyabovorn et al., 2023), failing to attract significant producer participation and hindering confidence in environmental and financial outcomes. In short, the identification and quantification of SOC change at a meaningful scale has been stymied by broad ranges and large uncertainties (Wiesmeier et al., 2019).

Individual process-based ecosystem models developed for agriculture, more precisely called cropping system models (CSMs) (Jones et al., 2017), have been used to investigate the effects of management and climate change on SOC and GHG emissions (Basso et al., 2018; Campbell and Paustian, 2015; Grace et al., 2006) at regional to global scales (e.g., Del Grosso et al., 2006; Grace and Robertson, 2021). CSMs incorporate varying theoretical assumptions, often using limited empirical observations for model calibration and validation (Sulman et al., 2018; Wallach et al., 2021). Owing to the different crops, management practices, and geographic regions for which they have been developed, different CSMs can provide divergent predictions for SOC change, particularly for those models applied to environments and scenarios for which they have not been calibrated or developed (Rosenzweig et al., 2014). Unsurprising, given that calibration is challenging, often unfeasible at large scale, lacking independent validation (Le Noë et al., 2023), and with model parametrization that contains bias leading to estimation errors (Gaillard et al., 2018; Wallach et al., 2021) and unsuitable business-as-usual (BAU) baselines (Olson, 2013).

Research over the last several decades has shown that there is no silver-bullet SOC model (Paustian et al., 2016), although some have been more widely tested than others, and some specific combinations of models can perform better (Riggers et al., 2019). So, even if CSMs provide an attractive alternative to large-scale SOC measurement programs, no one model can satisfactorily quantify its change across all combinations of soils, climates, crops, and agronomic practices at varying scales, despite the well-validated performance of individual models in specific contexts. For all these reasons, models practitioners and certificate providers still face the challenge of selecting the right model, appropriate to the specific conditions in which they are working (Garsia et al., 2023).

Climate modelers have addressed this problem with multi-model ensemble (MME) approaches (e.g., Basso et al., 2018), which have also worked well for predicting climate-induced crop productivity change (Jägermeyr et al., 2021; Martre et al., 2015). MMEs capture differences in model structure as well as broader sets of environmental conditions and management practices than those for which individual models have been developed and tested. Consequently, they provide a substantial advantage for evaluating SOC change in response to changes in management practices and climate, and could be particularly attractive for addressing measurement, monitoring, reporting, and verification (MMRV) challenges of current carbon credit frameworks (Oldfield et al., 2022; Wongpiyabovorn et al., 2023). At small scales, previous MMEs have been used to predict SOC change more accurately than individual models in croplands (e.g. Basso et al., 2018; Riggers et al., 2019; Sándor et al., 2020), grasslands (e.g., Fuchs et al., 2020), and bare soil (e.g., Farina et al., 2021), but have yet to be applied to large, landscape-scale questions despite the scientific and policy importance of mitigation interventions in agriculture (Lecocq et al., 2022).

Soil sampling programs typically cannot meet MMRV challenges because of the cost of sampling and analysis, the need to adequately capture spatial variability, and the decadal or longer time frame required to detect credible SOC change (Brinton et al., 2025; Even et al., 2025; Goidts et al., 2009; Stanley et al., 2023). Consequently, on-farm observations, unless conducted as part of funded scientific trials with scientific expert supervision, typically cannot reliably capture the magnitude of carbon dynamics, particularly the rate of SOC change (Bradford et al., 2023), leading to large uncertainties in SOC datasets (Potash et al., 2022). These are major problems for calculating cropland soil carbon budgets and credits (Zhou et al., 2023), with important implications for crediting soil carbon gains (Oldfield et al., 2022).

Here we developed an MME comprised of widely used CSMs (APSIM, ARMOSA, CropSyst, DayCent, DSSAT, EPIC, SALUS, and STICS) to (1) evaluate its performance against individual-model simulations results over long-term research sites with field observations; (2) estimate changes in SOC stocks for conventional versus regenerative cropland practices at 40,000 unique locations across 934 counties in 12 US Midwest States.

## 2. Material and Methods

### 2.1 Process-based Cropping Systems Models (CSMs)

Eight different process-based CSMs were used to develop the MME (Table S1): Apsim (Keating et al., 2003), Armosa (Perego et al., 2013), Cropsyst (Stöckle et al., 2003), Daycent (Parton et al., 1998), Dssat (Jones et al., 2003), Epic (Izaurralde et al., 2006), Salus (Basso and Ritchie, 2015) and Stics (Beaudoin et al., 2023). Models were selected primarily based on their ability to reproduce interactions between soil, plant, and the atmosphere to simulate crop yields and ecosystem processes, including SOC change. Each model uniquely simulates and mathematically represents these processes (Table S2) and has been shown to appropriately represent SOC change under well-validated field conditions representative of cropping systems and soils of the US Midwest cropland. All models operate with daily weather inputs and can simulate carbon and nitrogen cycling, SOC change, nitrous oxide emissions, water balance, and plant growth in response to different crop management practices including fertilization, tillage, and irrigation. A brief description of each model is provided in the supplemental material (Section 1).

### 2.2 Scaling the Multi-Model Ensemble

The MME was scaled to cover the US Midwest cropland over a total of ∼46 million hectares that includes large areas in the 12 states of Illinois, Indiana, Iowa, Kansas, Michigan, Minnesota, Missouri, Nebraska, North Dakota, Ohio, South Dakota, and Wisconsin. Simulations were performed over 42 years (1980-2022) using gridMET weather observations data (see 2.2.2).

#### 2.2.1 Management scenarios and dynamic baselines

Eight management scenarios were simulated to evaluate SOC dynamics (Table 1), four (1-4) with full nitrogen (FN) fertilization and four (5-8) with reduced N fertilization (RN, Table S3). These scenarios constitute business-as-usual (BAU) and regenerative agricultural practices and crop rotations across the US Midwest, reflecting those most frequently studied in the literature at test site locations (see 2.3.1). Crop rotations include maize and soybean with or without a cover crop (Table S4) and with either conventional tillage or continuous no-till (Table S5). These scenarios allow comparisons of SOC change due to different management practices over the long-term at specific locations and at multiple scales using a dynamic baseline, a time and space dependent reference that accounts for natural and management-driven variations in SOC levels.

**Table 1.**
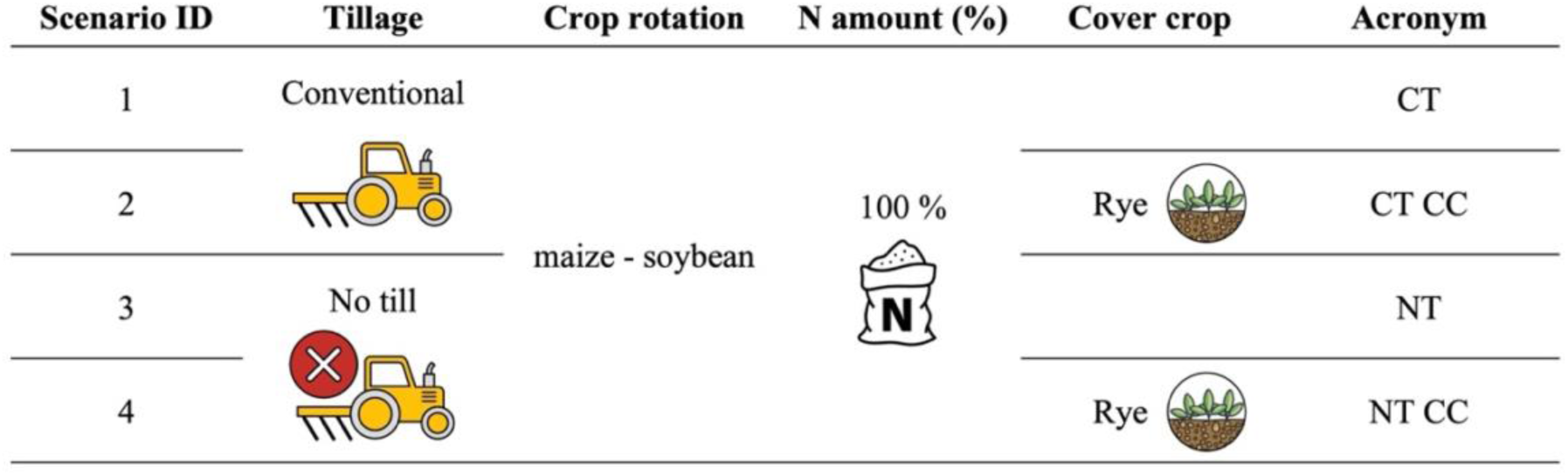
Scenarios simulated using a multi-model ensemble (MME) approach, representing different combinations of management practices involving tillage and cover cropping. The full nitrogen fertilization (100 %) refers to the Midwest state based “business as usual” average amount and was applied only for maize. A two-year crop rotation, with maize followed by soybean, was implemented in all scenarios. Scenario acronyms: CT (conventional till), NT (no till), FN (full nitrogen), CC (cover crop). Graphical icons illustrate the different practices and are adopted throughout the figures.

| Scenario ID | Tillage | Crop rotation | N amount (%) | Cover crop | Acronym |
| --- | --- | --- | --- | --- | --- |
| 1 | Conventional | maize - soybean | 100 % |  | CT |
| 2 |  |  |  | Rye | CT CC |
| 3 | No till |  |  |  | NT |
| 4 |  |  |  | Rye | NT CC |

#### 2.2.2 Data inputs

Simulations were conducted at 1/24^th^ degree spatial resolution as determined by gridMET weather observations (Abatzoglou, 2013). Each grid (approximately 4 km^2^) represents an individual simulation-based unit (with a geolocated centroid) labeled with a unique identifier (UID). Grids were extracted for the 12 Midwest states involved in the analysis. The cropland area (maize or soybean grown for at least one year between 2008 to 2022) within each grid was calculated, as provided (30 m resolution) by the USDA National Agricultural Statistics Service Cropland Data Layer (NASS, 2024). The percentage of cropland area per grid was calculated by dividing the cropland area by the total cropland area summed across the 12 Midwest states. Then with grids ordered from smallest to largest percent cropland, the percentages were summed to calculate the cumulative percent of cropland area. We selected UIDs that when combined covered 90% of the cropland area (Figure S1). Selected UIDs were then categorized into discrete latitude bins to enable more straightforward identification and homogenous extraction of the final number of UIDs to be run. Within each bin, the main soil textures were identified, and their total area (km^2^) determined. Forty thousand total UIDs (balancing the computational power required and area representability) were then chosen across the region using a weighted random selection process, where the number of grids selected within each latitude and bin-texture combination was proportional to the area of cropland within the bin and the area covered by each texture. This selection process gave a higher probability for grids with a larger crop area to be selected.

For each UID, daily variables of maximum temperature, minimum temperature, precipitation accumulation, downward surface shortwave radiation, wind velocity, maximum relative humidity, and minimum relative humidity were downloaded from gridMET for 1980 to 2022.

Soil data for model inputs were taken from the USDA Natural Resources Conservation Service Gridded Soil Survey Geographic (gSSURGO) (Soil Survey Staff, 2020) at 30 m spatial resolution. Variables extracted included bulk density, sand/silt/clay content, stone content, organic matter, and pH for each layer in the soil profile, using the most representative component percentage from the gSSURGO database. From these data, the following variables were calculated: organic carbon, total nitrogen, drained upper limit and lower limit (Ritchie et al., 1999), and saturated hydraulic conductivity (Suleiman and Ritchie, 2001). The dominant soil type within each weather grid was selected to represent that specific weather grid, excluding missing or shallower than 30 cm soil profiles.

Annual planting and harvesting dates for maize and soybean for 1980-2022 were estimated from USDA NASS state-level weekly progress reports, spatially interpolated with county-level annual crop production area. Daily progress of planting or harvesting was linearly interpolated between the weekly progress reports to find when 50% of the state was planted or harvested.

State centroids were computed, weighted by county-level crop production. The planting or harvesting dates were spatially interpolated between the weighted state centroids using a generalized additive model (GAM). The model is defined as:

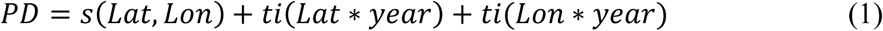

where *PD* is the predicted date, *s(Lat, Lon)* is the two-dimensional smoothing function with latitude and longitude as covariates, and *ti(Lat*year)* and *ti(Lat*year)* are tensor functions of the random effect of year and latitude and longitude, respectively. The model was applied to the weather grid centroids for each year, predicting planting and harvesting dates for each weather grid. Missing annual county-level production data was filled using the average across all available years. When all data was missing (e.g., soybean progress reports for 1979), the long-term average centroid and progress values were used in the model.

Nitrogen (N) fertilizer application amounts for maize were based on the USDA ARMS survey as reported in (Basso, 2019). The average N rate was used to represent each state, except for Wisconsin where the maximum reported N rate was used to account for the application of manure not included in the reported range of N rates, as indicated by the relatively low range, and for Kansas and Nebraska, where data were not available, and the same rate as a nearby state (Missouri and Iowa, respectively) were applied (Table S6).

#### 2.2.3 Crop maturity regions identification and cultivars parametrization

Crop maturity regions (MR) were identified to better represent crop cultivar extent across the US Midwest. These regions were based on six latitude bins each for maize and soybean, with each bin determined by the crop growing degree days (GDD) accumulated at maturity. Initially, the GDD data for maize for each UID were derived from (Abendroth et al., 2021) and for soybean, computed from weather data (according to NASS planting and harvest dates). Then, the distinctions between each MR for each crop were computed by calculating the quantile-based breaks for the desired number of bins (6). These breaks were then used to categorize all UIDs into a distinct maturity region group (Figure S2). The final GDD values for each MR was computed as the median of each group (Table S7).

For each CSM, crop cultivars were then parametrized using the GDD value according to the MR. In addition, to ensure consistency across CSM, harvest index (HI) and crop base temperature were set to standard values. Defaults for HI were initially set to 0.5 and 0.4 for maize and soybean, respectively (some CSMs then recalculated these HI values throughout the simulation period), and for crop base temperature were 8°C and 10°C for maize and soybean, respectively. Thus, this rationale was intended to set common main crop parameters across models to simulate crop yield (Figures S3 and S4).

#### 2.2.4 Multi-Model Ensemble computation

Daily simulation using the eight CSMs over 42 years across 40,000 UIDs in the US Midwest cropland generated a large dataset of more than 4 billion data values per output metric. For each model and UID, the standardized output data was aggregated first by year and then grouped into a unique dataset. Then, for each management scenario, the yearly data were grouped by county. Outliers were detected using the interquartile range (IQR) method. This function calculates the range between the first (25th percentile) and third (75th percentile) quartile and defines outliers as data points that fall below the first quartile minus 1.5 times the IQR or above the third quartile plus 1.5 times the IQR. Once outliers were removed, the MME median (e-median) was used as the estimator of the ensemble model simulations.

When aggregating data by county, a weighted mean was used to account for the varying cropland area extents associated with each UID within the county.

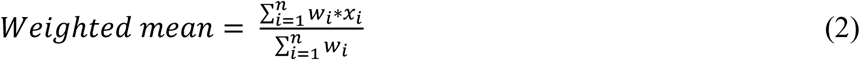

where *x*_*i*_ is the value of the i*^th^* UID value and *W*_*i*_ is the weight associated with the i*^th^* observation. Each weight is assigned based on the cropland area of each UID (i.e., grid). Equal weight was adopted across models. Scenario outputs in terms of delta SOC change of N2O emissions are reported throughout the manuscript as mean and standard deviation at the county level. The delta SOC change for the MME was calculated as a mean of each annual SOC change. R software was used for statistical analysis and graphical visualization.

### 2.3 Multi-Model Ensemble validation

To test the MME performance in reproducing SOC change, we evaluated its accuracy and uncertainty in comparison with every individual CSMs. The MME and each CSM were evaluated comparing the simulated SOC stock and annual SOC change values for different management practices (i.e., experimental treatments) with observed (measured) values from 17 experimental sites (see 2.3.1). We tested MME and CSMs without performing a specific calibration for the soil-related parameters. Our goal was to demonstrate that the MME is suitable for upscaling over large areas where site specific soil parameter calibration is not feasible.

Therefore, we rely primarily on the correct crop yield settings for the upscale and experimental sites. Lastly, MME uncertainties were also evaluated on the upscaled simulations across the entire US Midwest.

#### 2.3.1 Experimental sites with observed data

To ensure consistent, high quality observed data sets were used for individual CSMs yield calibration and MME and CSMs validation, experimental sites were only selected from the literature when five data conditions were present: [1] initial soil conditions (prior to experimental treatments), including SOC stock, [2] multiple years of crop yield data, [3] a SOC time series’ (with either SOC stock values or BD and C% available for SOC stock calculation), [4] detailed management descriptions and ancillary information, and [5] the presence of maize or soybean, grown either continuously, or in a two-year rotation, or within an extended rotation with other crops (e.g., wheat and alfalfa). Sites were also intentionally selected to cover a wide range of management, including conventional tillage, reduced tillage, no-till, different fertilization rates, and the presence of cover crops. Sampling depth up to 30 cm was preferred, but in 10 locations the depth of sampling was less (i.e., between 10 and 25 cm). In total, 17 experimental locations, with 52 site treatments, were located in North America (16 in the USA and 1 in Canada) (Figure S5).

To simulate these sites, weather data was extracted from gridMET for US sites and NASA Power (NASA Langley Research Center, 2024) for Canadian sites. Soil data were preferentially and primarily sourced from the publications, but if unavailable, gSSURGO data at the closest site were used. Water limit properties and saturation were recalculated using pedotransfer functions as reported above. If not reported, initial soil water content was set equal to the drained upper limit (DUL) and initial inorganic nitrogen was set to 40 kg N ha^-1^ for the entire soil profile depth, with ammonium and nitrate ratios assigned based on each model’s default. Table S8 reports a full description of all sites and treatments.

To ensure appropriate biomass feedback from the crop residues and roots, each CSM was calibrated at each experimental site only against the observed crop yield data without involving any soil-related parameters. All the selected CSMs are designed to accurately predict crop yields, root and residue biomass (includes total above-ground biomass minus the grain yield) because of their critical importance for modeling long-term SOC dynamics. The RRMSE (Relative Root Mean Square Error, equation 3) was used to evaluate individual CSMs calibration performance (Table S9). Most of the time, total biomass of cover crop was not reported, and default models’ values were used to set its productivity. Data_S1, in the supplemental material, reports for each CSM and location, with the full set of parameters used, their default values, and the final values obtained from the calibration process.

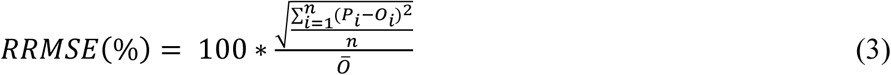

where *P*_*i*_ and *O*_*i*_ are predicted and observed data respectively, *n* the number of P/O pairs and *O̅* the mean of observed data.

#### 2.3.2 Multi-Model Ensemble and individual model accuracy and uncertainty analysis

Before evaluating the MME and each CSM’s performance in terms of accuracy and uncertainty, we compared the measured delta SOC change values from the 52 experimental site treatments with a recent meta-analysis on regenerative practices (Bai et al., 2019) to identify values that were inconsistent with the literature range, i.e., for no-till (-0.5-1.22 Mg ha^-1^ yr^-1^), conventional tillage (-0.47-0.5 Mg ha^-1^ yr^-1^), and reduced tillage (-0.97-0.65 Mg ha^-1^ yr^-1^). To do so, we calculated the weighted mean delta SOC change for the measured data (equation 2; where *x*_*i*_ is the i*^th^* measured delta SOC change and *W*_*i*_ its weight based on time elapsed since the previous SOC observation), and noted for each site*treatment whether it was consistent with the data range (Table S10). For the accuracy and uncertainty analyses described below, individual paired comparisons between simulated and observed data were included only when observed weighted mean annual delta SOC change were consistent with the literature ranges reported above. For the MME and individual CSMs instead, the weighted mean annual delta SOC change for was calculated as the average of the annual SOC changes over the simulated time period, with weights applied to account for shorter durations in the initial or final years of the simulation.

To evaluate MME and CSM accuracy, a composite score was developed by combining two metrics: the Savage score (Savage, 1954), and the weighted standard deviation (*s*) of model ranks. The absolute value of the prediction errors (PE, equation 4), computed as the difference between predicted (i.e., simulated) and measured delta SOC change were used to rank every j*^th^* model accuracy within each i*^th^* site*treatment combination.

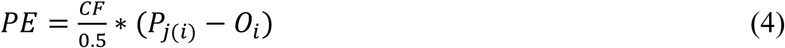

where *Pj(i)* and *Oi* are the predicted and measured values of the j*^th^* model and i*^th^* site*treatments combination respectively, and CF is a correction factor that considers measurement uncertainties (Daren et al. 2007; Riggers et al., 2019).

Each site*treatment combination ranking was then transformed into a Savage score. The total Savage score (Si) for each model, calculated as the sum of its scores across all site*treatment combinations, was used to rank model accuracy, with greater weight assigned to models demonstrating consistently high performance. Rank variability, represented by the weighted standard deviation of ranks (*srank*), quantified the stability of model rankings. The weight (1/rank) of the standard deviation was included to favor models that consistently fluctuate among top-ranking positions, rather than those that exhibit variability in lower-ranking positions. Both metrics were normalized to a 0-1 scale and, for each model, the composite score (CS) was computed as:

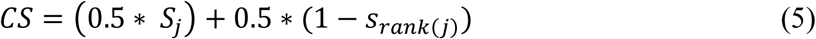

Where *Sj* is the normalized sum of the Savage-scores assigned to the j*^th^*model and *srank(j)* the weighted standard deviation of the j*^th^* model rank, while 0.5 assure the same weight to both metrics. This approach ensured that the highest-ranked models demonstrated both superior accuracy and low variability.

To assess uncertainty, we used the mean squared error of prediction (MSEP) and its decomposition in prediction squared bias (*bias*^2^) and prediction variance (*var*) as reported by Maiorano et al. (2017) on the raw SOC stock variable. This metric was used to quantify MME uncertainty in comparison with individual CSMs. Since no calibration was performed against SOC data, measured data from experimental sites were used as the prediction dataset.

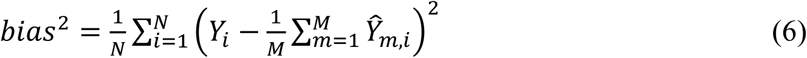

where *N* is the number of treatments, *Y*_*i*_ is the measured variable for the treatment *i*, *Ŷ*_*m,i*_ is the simulated variable for the treatment *i* by the *m* CSM (or MME). The predicted variance is the variance of the values simulated by the population of *m* models (1 when individual CSMs are tested) averaged across treatments:

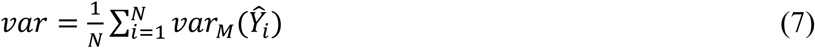

where *N* is the number of treatments, *Y*^_*i*_ is the simulated variable for the treatment i. The estimate of MSEP can then be obtained as:

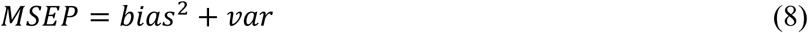

We further evaluated how the uncertainty estimates changed with the number of models involved using the coefficient of variation of the multi-model ensemble e-median over the upscaled dataset in the US Midwest. To do so, we performed a bootstrap calculation (i.e. random sampling with replacement) for each value of *M* (number of models in the ensemble) from 1 to 8. For each ensemble size *M* we extracted 10000 bootstrap samples of *M* models with replacement. This bootstrap sampling was repeated *N* times equal to the number of scenarios (1-4), and *C* times equal to the number of counties involved (934), considering single UID 40-years overall *C* gain/loss values within each county. The total number of samplings with replacement exceeded 35 million. The variation of the e-median across the bootstrap samples due to random model selection was computed with the coefficient of variation (CV):

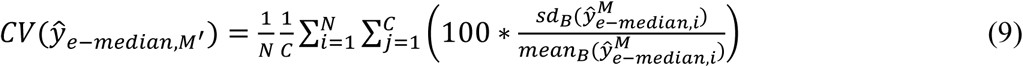

where *CV*(*ŷ*_*e*−*median,M*′_) is the coefficient of variation of e-median for the model ensemble of size *M*, *sd*_*B*_ and *mean*_*B*_ are the standard deviation and the mean of the *B* (number of bootstrap sample) e-medians of the model ensemble of size *M* for the i*^th^* scenario (from 1 to 4) and the j*^th^* county.

## 3. Results

### 3.1 Comparison of Multi-Model Ensemble and Cropping System Models

In comparison with individual CSMs, MME best simulates changes in SOC across the experimental sites and treatments, with the highest overall composite score of 0.948 (Table 2). The MME Savage score of 39.3, indicates that it frequently ranks among the top models in terms of accuracy (i.e., lowest absolute prediction error) and also has the lowest weighted standard deviation of the absolute prediction error rank (2.07; Table 2). Uncertainty assessment using MSEP and its decomposition into squared bias and variance components, shows MME consistently exhibits the lowest uncertainty, evident when compared with the full CSM population (Figure S6) and with individual CSMs (Figure S7). When upscaled to the US Midwest, the coefficient of variation for the delta SOC change e-median decreases as the number of models increase, from 99% with one CSM to 36% with eight CSMs (Figure S8).

**Table 2.** Comparison of the performance of individual CSMs and the MME across measured experimental sites and treatments. The Savage score was calculated by ranking the prediction errors between measured and simulated delta SOC changes within each site and treatment combination (Table S8). The weighted standard deviation reflects the variability in performance rankings across locations and treatments for each model with respect of the top rank position compared to the bottom ones. The composite score represents the overall performance, assigning greater weight to higher Savage scores and lower standard deviations, and is calculated using Equation 5. Higher composite scores indicate better overall performance. Normalized values are presented within parenthesis for the Savage score and weighted standard deviation respectively.

| **Overall**<br>**performance**<br>**ranking** | **Model ID** | **Savage score of**<br>**the absolute**<br>**prediction error rank** | **Weighted**<br>**standard deviation of the**<br>**absolute prediction error rank** | **Composite**<br>**performance score** |
| --- | --- | --- | --- | --- |
| **1** | MME | 39.3 (0.90) | 2.07 (0.00) | 0.948 |
| **2** | 3 | 41.5 (1.00) | 2.17 (0.11) | 0.946 |
| **3** | 7 | 37.3 (0.80) | 2.23 (0.17) | 0.813 |
| **4** | 6 | 34.9 (0.68) | 2.18 (0.12) | 0.781 |
| **5** | 2 | 32.9 (0.58) | 2.31 (0.26) | 0.663 |
| **6** | 5 | 28.2 (0.36) | 2.80 (0.79) | 0.285 |
| **7** | 1 | 27.5 (0.32) | 2.80 (0.79) | 0.268 |
| **8** | 8 | 25.4 (0.22) | 3.00 (1.00) | 0.108 |
| **9** | 4 | 20.9 (0.00) | 2.81 (0.79) | 0.106 |

### 3.2 Regenerative practices and SOC change

Evaluation across ∼46.2 million hectares of US Midwest cropland using MME results grouped by counties, shows that in comparison with a dynamic baseline of conventional till without cover crops (CT), combined adoption of no-till and cover crops (NT CC; Scenario 4; Table 1) increases SOC accrual annually by 16.4 Tg C (on average 0.36 ± 0.12 Mg C ha^-1^ yr^-1^, Figure 1). This rate of increase is approximately twice that of when either no-till (NT; Scenario 3) or cover crops (CT CC; Scenario 2) are practiced independently on conventional till (i.e., 8.9 Tg C or 0.19 ± 0.13 Mg C ha^-1^ yr^-1^ for NT and 8.5 Tg C or 0.18 ± 0.14 Mg C ha^-1^ yr^-1^ for CC, Figure 2).

**Figure 1.**
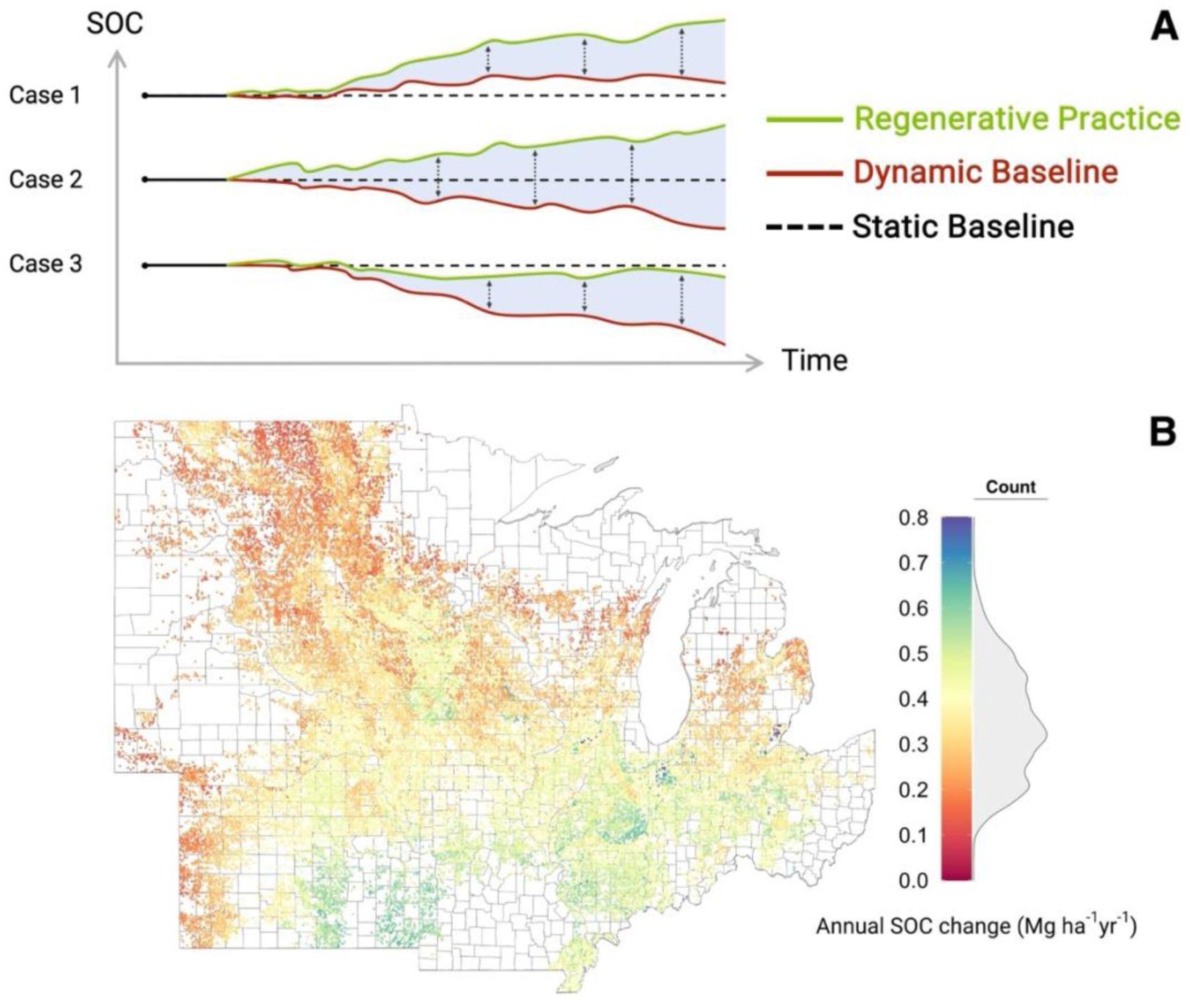
*Representation of the dynamic baseline concept and the mean MME annual SOC change difference between scenarios.* (A) The dynamic baselines (red), as well as the regenerative practice lines (green), account for yearly climate variability and site-specific soil properties, while the static baseline reflects the assumption of steady-state conditions. Three different cases are provided as examples of potential errors if changes are compared to static baseline and not the dynamic baseline; (B) MME annual soil organic carbon (SOC) change (Mg ha^-1^ yr^-1^, 0-30 cm) shown as the difference between Scenario 4 (NT FN CC, no-till with full nitrogen fertilization and cover crops) and Scenario 1 (CT FN, conventional till with full nitrogen fertilization) for each UID included in the study (n = 40,000).The gray density histograms represent the distribution of UID values across the US Midwest cropland.

**Figure 2.**
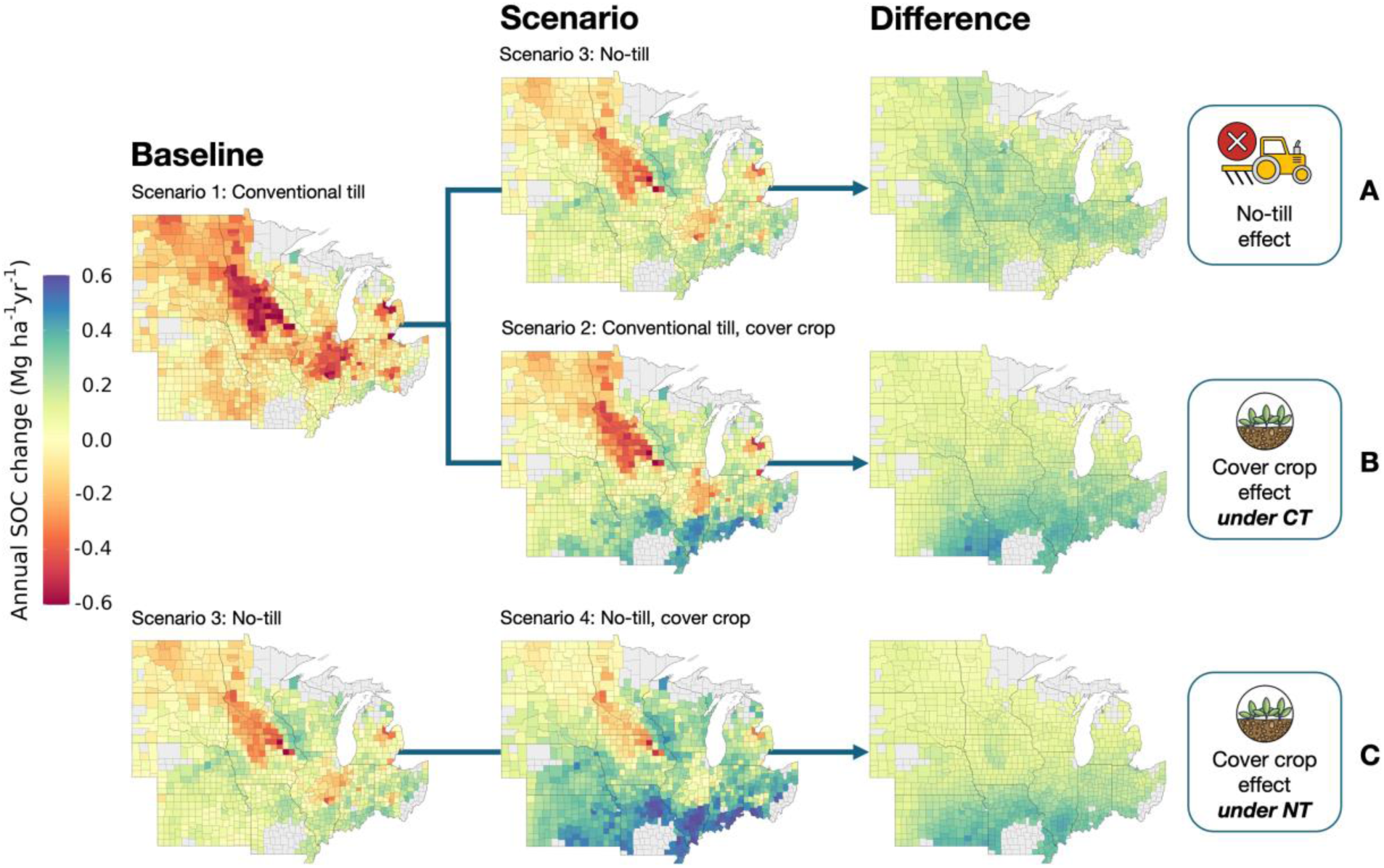
Effect of different regenerative practices on county-based MME mean annual SOC change. (**A**) Effect of no-till on county annual soil organic carbon (SOC) change (Mg ha^-1^ yr^-1^, 0-30 cm) shown as the difference between Scenario 3 (NT FN, no till with full nitrogen fertilization) and the baseline Scenario 1 (CT FN, conventional till and full nitrogen fertilization); (**B**) effect of cover crop on county annual soil organic carbon (SOC) change (Mg ha^-1^ yr^-1^, 0-30 cm) shown as the difference between Scenario 2 (CT FN CC, conventional till with full nitrogen fertilization and cover crop) and the baseline Scenario 1 (CT FN, conventional till and full nitrogen fertilization); (**C**) effect of cover crop under no-till on county annual soil organic carbon (SOC) change (Mg ha^-1^ yr^-1^, 0-30 cm) shown as the difference between Scenario 4 (NT FN CC, no till, full nitrogen fertilization and cover crop) and the baseline Scenario 3 (NT FN, no till with full nitrogen fertilization). Baselines serve as benchmarks for comparison with different scenarios. Individual counties are represented by their mean weighted values, with weights accounting for the agricultural area percentage of each county’s UID.

Similarly, under no-till, the introduction of a cover crop (NT CC) increases SOC accrual by 7.5 Tg C (on average 0.16 ± 0.10 Mg C ha^-1^ yr^-1^, Figure 2C). See Figures S9 and S10 for a full set of management comparisons. When N fertilizer is reduced to 75% of the BAU rate (Scenarios 5 vs. 1, 6 vs. 2, 7 vs. 3, and 8 vs. 4), SOC stock changes are negligible, averaging -0.02 Mg C ha^-1^ yr^-1^, while averaged across the RN scenarios N2O emissions decrease by 15% (-0.22 ± 0.45 kg ha^-1^ yr^-1^), and with cover crop adoption, increase by 11% (0.15 ± 0.51 kg ha^-1^ yr^-1^) and 12% (0.16 ± 0.45 kg ha^-1^ yr^-1^) under conventional and no-till management (Figure S11), respectively. See Figures S12 to S14 for a full set of county-level annual delta SOC and N2O emissions per scenario.

### 3.3 Pedo-climatic variables and SOC change

Using MME analysis, locations with higher initial carbon stocks (see Figure S15) typically show SOC loss (up to -0.43 Mg C ha^-1^ yr^-1^ in sandy soils under CT; Figure 3). Conversely, soils with low initial carbon stocks typically show SOC increases under both CT and NT, particularly with cover crops (CT CC and NT CC). Larger, consistent SOC accrual rates were apparent across soil textures and initial carbon stock when no-till and cover crops (NT CC) were present together, with similar trends emerging for RN scenarios (Figure S16). When individual CSMs were evaluated, changes in pedo-climatic variables inputs (initial SOC stock, clay content, and latitude; Figure 4) show no consistent impact on annual SOC change. As initial SOC stock increases, a decrease in the soil’s capacity to store more C is observed (CSMs 1, 3, 4, 5, and 6), but some show variable (CSMs 7 and 8) and no (CSM 2) response to initial SOC stocks.

**Figure 3.**
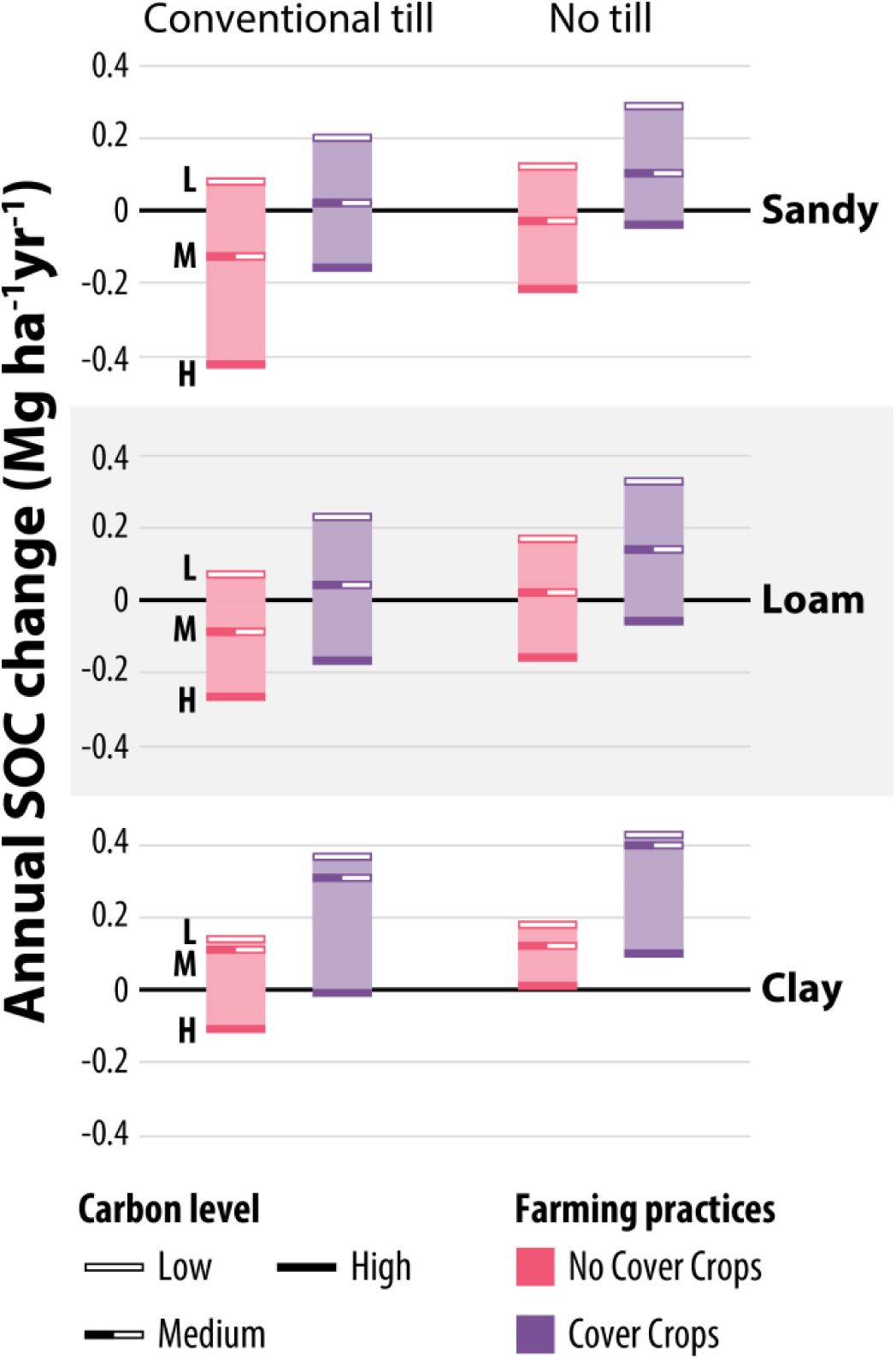
MME mean annual SOC change rates across scenarios based on different soil textures and initial SOC stock levels across 46 M ha. Annual soil organic carbon (SOC) change (Mg ha^-1^ yr^-1^, 0-30 cm) across scenarios (1 to 4, from left to right), soil texture (Sandy, Loam and Clay), and initial SOC stock level (Low, < 40 Mg C ha^-1^; Medium, 40- 80 Mg C ha^-1^; and High, > 80 Mg C ha^-1^). Annual SOC change values derived from all UIDs mean across the US Midwest cropland. All scenarios use maize-soybean crop rotation.

**Figure 4.**
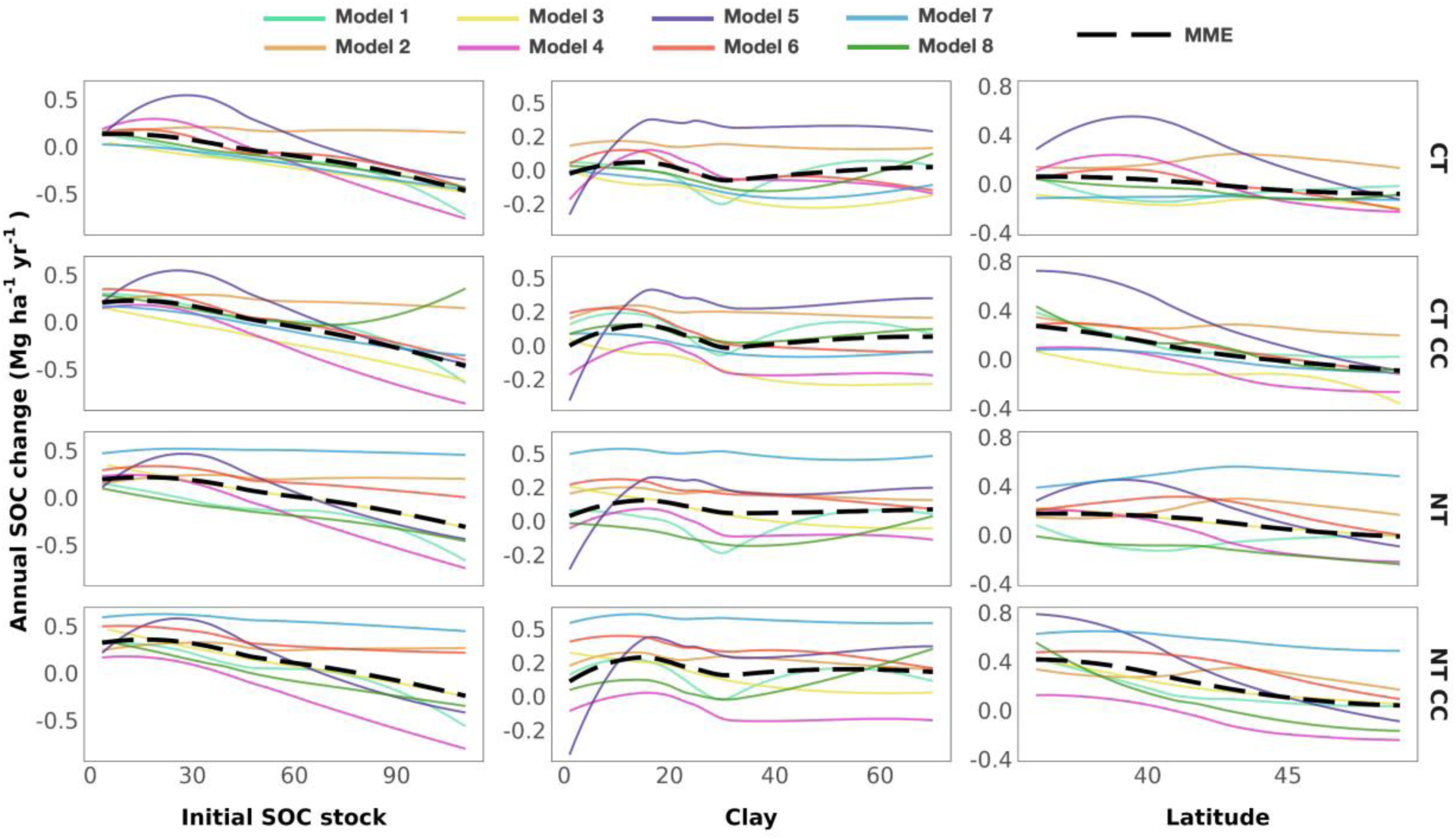
MME mean annual soil organic carbon change response to variations in initial SOC stock, clay content, and latitude across 46 M ha. Each line, represented by a unique color, corresponds to an individual model, with the black dotted line the MME median. They represent individual model annual soil organic carbon (SOC) change (Mg ha^-1^ yr^-1^, 0-15 cm) response to variations in initial SOC stock (Mg ha^-1^), clay content (%), and latitude (°). The different scenarios (1 to 4 from top to bottom) are displayed at the right end of each row. Scenario acronyms: CT (conventional till), NT (no till), FN (full nitrogen), CC (cover crop).

Increasing clay content beyond ∼30% shows an increase (CSMs 1 and 8), no change (CSMs 2, 4, and 7), or a decrease (CSM 3) in SOC stock. Soils at higher latitude show lower annual SOC change in CT scenarios but were inconsistent under NT management.

## 4. Discussion

### 4.1 Standardizing SOC quantification

Adopting regenerative management practices can reduce the carbon footprint of row-crop agriculture, but to what degree and whether it is meaningful as part of climate mitigation efforts is debated (Schlesinger, 2022). A major issue in determining impact is the lack of standardization in quantifying SOC stock change (Smith et al., 2020); estimates for individual fields based on re- inventorying, even after extended periods of time and at high sampling densities, can be variable and inaccurate, with gains or losses in SOC stock often determined even when no change has occurred (Bradford et al., 2023). These challenges are consistent with our findings from comparisons of SOC changes from individual long-term experimental treatments (Table S8) with meta-analysis ranges (Bai et al., 2019), where 38% (20 out of 52) of the peer reviewed data fell outside the range, some showing ‘unrealistic’ rates of change (e.g., 8 Mg C ha^-1^ yr^-1^; Table S10). The likelihood for spurious values is increased when SOC change is determined using consecutive short-term sampling (e.g., 4-5 years apart), typical with current C market approaches for paying producers, where contract lengths of only one year are common (Wongpiyabovorn et al., 2023). Short term contracts that eschew consideration of long-term SOC trajectories, risk exposure to measurements that are strongly impacted by seasonal trends (e.g., saw-tooth dynamics). While acknowledging high quality SOC field measurements are needed to better understand SOC dynamics and help to initialize models, their use in determining SOC stock change is problematic for short-duration payment requirements, and effectively unfeasible for larger-scale projects.

### 4.2 Multi-Model Ensemble performance

MME reduces the biases associated with short-term sampling and the use of individual CSMs, providing higher accuracy, and lower uncertainty (Table 2). An estimate from an individual CSMs that is ‘best’ at one field may not be so at another, often for reasons that cannot readily be determined; individual CSMs have varying constraints and needs and are better suited for specific contexts (Garsia et al., 2023). Notable disagreement among individual CSMs at large scale for the same scenarios (Figures S17–S21), underscores the quantitative lottery of individual CSMs selection for a particular application. Thus, MME provides a more consistent and reliable output of long-term SOC change across multiple scales in agreement with prior studies (Farina et al. 2021; Riggers et al., 2019).

### 4.3 Upscaling regenerative practices

By using high spatial resolution data aggregated to the US Midwest cropland, MME SOC change values are applicable and consistent with observations at multiple scales. As a management option, combining cover crops and no-till shows greatest promise (∼16.4 Tg C yr^-1^) averaged over the total area investigated. With cover crop introduction, the MME SOC increase of 0.17 Mg C ha^-1^ yr^-1^ (∼8.0 Tg C yr^-1^) across tillage types, is consistent with the meta-analyses of Peng et al. (2023) and McClelland et al. (2021) who found increases of 0.24 Mg C ha^-1^ yr^-1^ and 0.21 Mg C ha^-1^ yr^-1^, respectively. Over similar total areas of US cropland, potential SOC accrual suggests that MME provides a reasonable balanced estimate between non-conservative values that use IPCC factors (Sperow, 2020; 13.3 and 17.7 Tg C yr^-1^ for no-till and cover crop adoption, respectively), and conservative evaluation that uses satellite imagery and longitudinal surveys (Uludere Aragon et al. 2024; 5.3 Tg C yr^-1^ for cover crop adoption).

Although not our focus, quantifying N2O emissions is essential for estimating the broader global warming impact (GWI) of regenerative practices. Even with no validation, MME estimates align with literature values. For example, cover crop adoption increases N2O emissions by on average ∼0.15 kg ha^-1^ yr^-1^ across tillage managements (Figure S11), negligible increases consistent with the variable results reported by Basche et al. (2014). Prior multi-model studies have found significant uncertainty and divergence of N2O emissions across models and have adopted different approaches that omit agricultural management and crop growth (Tian et al., 2019). Although we make no claim, an MME approach that includes these factors may be well suited to ‘filter’ out data outliers, often reported in single-model studies due to the challenges of accurate simulation, especially on daily time-scales; further studies are needed to confirm this.

### 4.4 Dynamic baselines and flexible scales

Inconsistent and contradictory outputs highlight the ongoing risk of using a single CSM to provide the robust accounting needed to support agricultural carbon market projects across broad regions. A large cause of this incongruency is the known and somewhat intractable challenge of determining additionality through the choice of a suitable BAU baseline: the counterfactual scenario that would have occurred without management intervention. This choice can have a large influence on the degree of SOC change, and, by extension, on how many carbon credits are generated; if inaccurate, too many or too few may be issued, calling into question legitimacy (Paustian et al., 2019). Indeed, only C that is additionally stored in soils or that is additional compared to a BAU scenario can be relevant for climate change mitigation (Don et al., 2024).

Usually, projected baselines are a ‘best guess’ manufactured scenario that consider steady-state historical conditions for a specific area (Figure. 1A) without accounting for within season climate variability and site-specific soil properties. Attributing SOC change to management using a static baseline is problematic because it is prone to errors if SOC changes in the counterfactual BAU scenario are not quantified. MME avoids this issue by using dynamic baselines that are temporally and spatially adjusted to capture the impacts of fluctuating environmental variables. This allows quantification of the yearly SOC change difference between an adopted regenerative management practice and its dynamic counterfactual baseline to provide a more credible estimate of carbon gained or lost. Dynamic baseline scenarios were determined for every UID to allow for direct comparisons at the same location.

MMEs can explore large pedo-climatic variability with a regionally consistent framework. Units of scale are flexible – they can be an agroecological zone with similar soils, weather, and agricultural practices, such as USDA Major Land Resource Areas (USDA, 2022) or a political jurisdiction such as county or state (Schwartzman et al., 2021). Irrespective, estimates of SOC change can be quantified relative to a dynamic baseline for the scale chosen, and the results reported in readily accessible lookup tables, with the MMEs potential to generate and retrieve this information a valuable service, when reference values are needed for carbon credit payments or a regional policy document. As an example, the choice of county scale as the baseline jurisdictional unit allows their relatively uniform distribution across the US Midwest to capture soil and climate variability, and their administrative structure to provide a practical, credible framework for investment and implementation of regenerative practices for GHG abatement.

### 4.5 Opportunities for Multi-Model Ensemble application

As part of its effort to reduce GHG emissions, the US has promoted programs to enhance soil carbon sinks (UNFCCC, 2021). The expectation is that emissions reductions will occur through the adoption of regenerative practices on cropland (USDA, 2025), with market-based interventions best positioned to achieve this (USDA, 2023b). Voluntary offset and inset initiatives predominate in the US agricultural sector, however carbon credits generated by agriculture constitute only ∼1% of the total volume transacted (Haya et al., 2024). Despite very high rates of producer carbon market awareness, participation rates are very low (Urban and Cole, 2022), in large part a reflection of the barriers that currently hinder engagement, primarily high enrollment costs driven by quantification, recordkeeping, and data analysis, scale and aggregation limitations, and verification, which are insufficiently compensated for by C change payments (Parkhurst et al., 2023; USDA, 2023b). Cultural and social factors can also deter participation, including confusion about the assortment of available market programs (Brokish et al., 2023; Oldfield et al., 2022; Wongpiyabovorn et al., 2023).

Use of an MME accounting tool could alleviate many producer participation barriers, ease onerous MMRV requirements, provide confidence in SOC stock change quantification at different scales, and in doing so address several strategic US policy priorities (USDA, 2023a). Producers with eligible fields would not need to provide detailed agronomic data or information to participate, minimizing record keeping and auditing burdens, and potentially privacy concerns. In effect, an MME approach provides multi-scale, pre-run, practice-based, dynamic baselines for determining emission factors with low uncertainty levels. MMEs may be a practical, intermediate approach for payment for practice and payment for output mechanisms. As a decision support tool, MMEs can also allow producers or project developers to quickly screen lands targeted for regenerative practices to determine their economic and program viability (USDA, 2023b).

Of the few registered agricultural land management (ALM) projects, most have generated low density (per hectare) and low volume (i.e., small area) credit numbers when compared to other sector project types (Haya et al., 2024; Smith and Parkhurst, 2018). Unless aggregation (combining individual fields or small projects) to form large scale projects is allowed, producers, particularly those with limited resources or opportunity, find cost barriers insurmountable. While aggregation in varied design is allowed in current agricultural protocols and results in verification cost reduction, it does not reduce quantification costs, which scale with the area enrolled.

Similarly, approaches and accounting rules have been recommended to address issues of permanence, leakage, and additionality, inherent with small-scale, individual ALM projects (Smith and Parkhurst, 2018). However aggregated projects where management interventions are accounted for at larger scales reduce these concerns (Santilli et al., 2005). With sufficiently large spatial and temporal scales, additionality can be demonstrated through reduced emissions below recent regional baseline trends. Larger project area coverage and longer project times can alleviate practice reversal risks, unintentional or otherwise, by pooling across broader scales and stakeholders. Large programs can also mitigate leakage risks by addressing underlying economic drivers of emissions and can reduce jurisdictional risk by directly accounting for all emissions shifts inside the administrative unit (Schwartzman, 2021). Measuring BAU baselines on a project-by-project basis is not economically feasible as emissions reductions are too small relative to costs. Technology-adjusted historical baselines designed in accordance with regional dynamics seem a superior option (Mendelsohn et al., 2021), and one that MMEs are well-suited to provide.

## Conclusions

MME results generate low uncertainty estimates of SOC stock change for common management practices that can be used in C market accounting at multiple scales. This removes the exposure of market actors to accusations of model selection and reduces the uncertainty of practice change outcomes. With MMEs, the fine granularity and consistent framework of input data across a large area allows for a flexible choice of scale for model output aggregation. This and the availability of practice-based dynamic baselines, along with decision support capabilities, can allow MMEs to reduce barriers to carbon market participation, and ameliorate long standing concerns related to additionality, leakage, and permanence. In doing so, MME addresses several policy priorities and has a potential role in national regenerative agriculture programs.

## Acknowledgments

We thank Roger L. Nelson for help with CropSyst; Isaiah L. Huber for help with APSIM, Marco Botta for help with ARMOSA, and Matt Wisniewski for help with figures.

## Funding

Partial funding to Basso is provided by: Great Lakes Bioenergy Research Center, U.S. Department of Energy, Office of Science, Biological and Environmental Research Program under Award Number DE-SC0018409; the National Science Foundation Long-term Ecological Research Program (DEB 2224712) at the Kellogg Biological Station, USDA NIFA, Award no. 2020-67021-32799, Michigan State University, AgBioResearch, Climate TRACE, CERCA-FFAR project, Walton Family Foundation, United Soybean Board. M. Delandmeter was granted a Research Fellow (number 44221) fellowship by the F.R.S.-FNRS (Belgian Fund for Scientific Research).

## Conflict of interest statement

Bruno Basso is a cofounder of CIBO Technologies. Keith Paustian and Yao Zhang has financial interest in Indigo Ag. The other coauthors declare no conflict of interest.

## Data availability statement

The data that support the findings of this study are openly available as supplemental material.

## Supporting Information

### 1. Single model descriptions

#### APSIM

The Agricultural Production Systems sIMulator (APSIM) is a model designed to simulate biophysical processes in agricultural systems, focusing on the economic and ecological outcomes of management practices under climate risk. It is also employed to explore solutions for food security, climate change adaptation and mitigation, and carbon trading. APSIM is organized into plant, soil, and management modules, which cover a wide range of crops, pastures, and trees, as well as soil processes such as water balance, nitrogen and phosphorus transformations, soil pH, erosion, and various management controls. APSIM was developed to meet the need for tools that provide accurate predictions of crop production in relation to climate, genotype, soil, and management factors while addressing long-term resource management issues.

#### ARMOSA

The ARMOSA model represents carbon and nitrogen fluxes and the influence of high-level agroecosystem processes that vary in response to agricultural management (i.e. crop rotation, crop residues management, fertilization, irrigation, tillage) and pedoclimatic conditions. At regional to national scale, it has the capability of depicting multi-crop rotations over a medium to long term perspective allowing quantification of crop production and environmental variables in response to varying market and policy needs (e.g., organic farming, greening, and healthy diet habits). The model runs at daily time step and consists of of three main modules: crop growth and development, soil water dynamics and C and N cycling.

#### CROPSYST

CropSyst is a multi-year, multi-crop, daily/hourly time step cropping systems simulation model developed to serve as an analytical tool to study the effects of climate, soils, and management on cropping systems. CropSyst simulates soil water budgets, nutrient budgets (N and P), C cycling, crop growth and yield, residue production, soil erosion, and other parameters under user-defined management options (including rotations, tillage, fertilization, and irrigation scheduling and options to deal with water shortages). CropSyst can provide insight into N pollution, GHG emissions, and C sequestration.

#### DAYCENT

DayCent is an ecosystem model designed to simulate terrestrial croplands, forest lands, grasslands, and savannas. The model predicts a variety of ecosystem variables including plant productivity, water balance, soil carbon, and nitrogen dynamic. DayCent is the daily time-step version of the Century model. DayCent consists of several sub-models, and it has been widely used and tested to quantify GHG fluxes (i.e., nitrous oxide and methane emissions) from agricultural soils for the U.S. National Greenhouse Gas Inventory. The model requires basic information on soil physical properties, plant type, and management practices as inputs.

#### DSSAT

The Decision Support System for Agrotechnology Transfer (DSSAT) is a software application that includes crop simulation models for more than 40 different crops. The software provides tools for efficient use of these models, such as database management programs for soil, weather, crop management, experimental data, and utilities and applications. The model simulates crop growth, development, and yield on a daily basis for a uniform area of land under specific management practices. Additionally, it simulates changes in soil water, carbon, and nitrogen levels over time within the cropping system.

#### EPIC

The Environmental Policy Integrated Climate (EPIC) model is a process-based computer model that simulates the physicochemical processes that occur in an agroecosystem under agricultural management. Management capabilities include irrigation, drainage, furrow diking, buffer strips, terraces, waterways, fertilization, manure management, lagoons, reservoirs, crop rotation, intercropping, pesticide application, grazing, and tillage. Besides these farm management functions, EPIC can be used in evaluating the effects of global climate/CO2 changes; designing environmentally safe, economic landfill sites; designing biomass production systems for energy; and other spin off applications. The major components in EPIC are weather simulation, hydrology, erosion-sedimentation, nutrient cycling, pesticide fate, plant growth, soil temperature, tillage, economics, and plant environment control.

#### SALUS

The SALUS (System Approach to Land Use Sustainability) program models a continuous cropping system under various management strategies for multiple years. These strategies may have various crop rotations, planting, irrigation, fertilizer, and tillage regimes. The program will simulate plant growth and soil conditions daily.

For each day and each management strategy, components of the crop-soil-water model are executed. These components are management practices, water balance, soil organic matter, nitrogen and phosphorous dynamics, heat balance, plant growth and plant development. The water balance considers runoff, infiltration, evaporation, soil water flow, drainage, root uptake, and transpiration. The soil model simulates organic matter decomposition, N mineralization and formation of ammonium and nitrate, N immobilization, and gaseous N. The development and growth of plants considers the environmental conditions (particularly temperature and light) to calculate the potential rates of growth for the plant. This growth is reduced based on water and nitrogen limitations.

#### STICS

The STICS (*Simulateur mulTIdisciplinaire pour les Cultures Standard* in French) model is a process-based soil-crop model which simulates plant growth as well as water, C and N fluxes. It runs at a daily time step and requires inputs related to weather, soil, crop and agricultural management, and provides outputs related to both agronomic (e.g. biomass and yield) and environmental (e.g. soil organic carbon, nitrate leaching, soil water and nitrogen, etc.) variables. STICS allows to investigate the impacts of contrasted pedo-climatic conditions and/or agricultural management (nitrogen fertilization, tillage, cover crops, etc.) on various crop rotations which might include grasslands. Each crop is represented as an ensemble of components whose variables (development stage, biomass, N content) evolve according to ecological functions. Using a 1D description, the model simulates the carbon, nitrogen and water fluxes over the soil profile and the whole plant cycle.

#### 2. SUPPLEMENTAL FIGURES

**Fig. S1.**
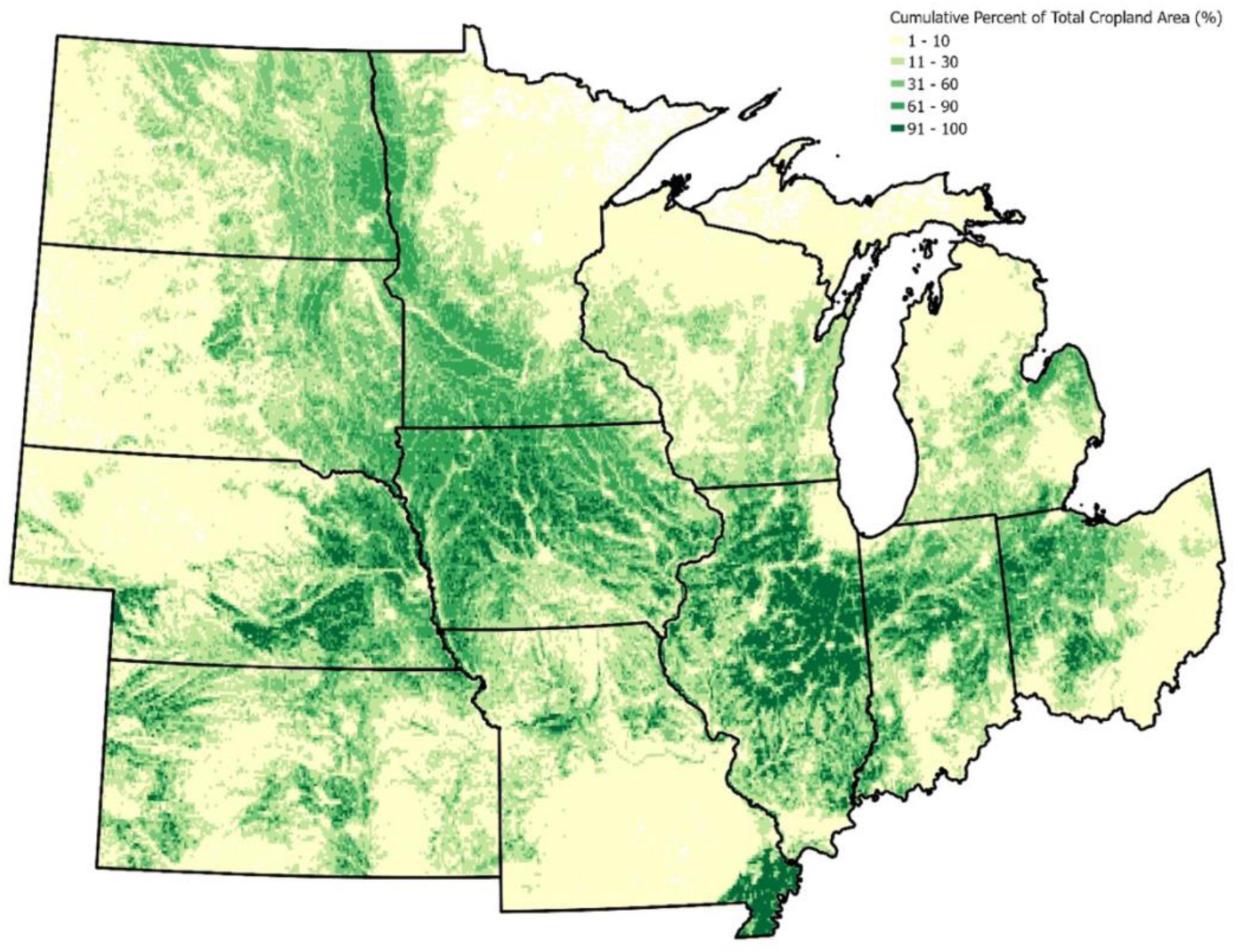
Cumulative percent (%) of total cropland area across gridMET weather grid cells.

**Fig. S2.**
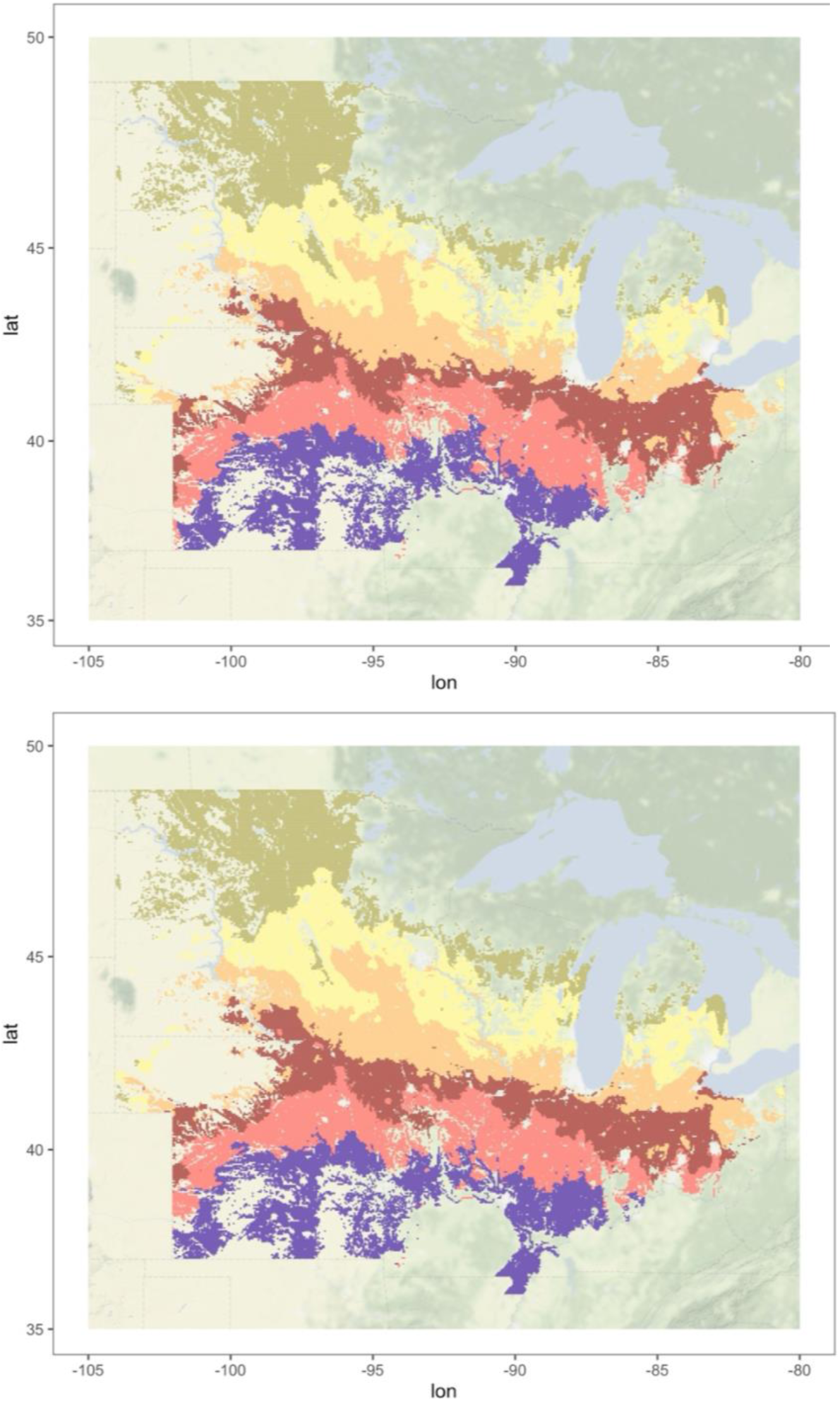
Distribution of the six maturity regions for soybean (top) and maize (bottom) across the US Midwest.

**Fig. S3.**
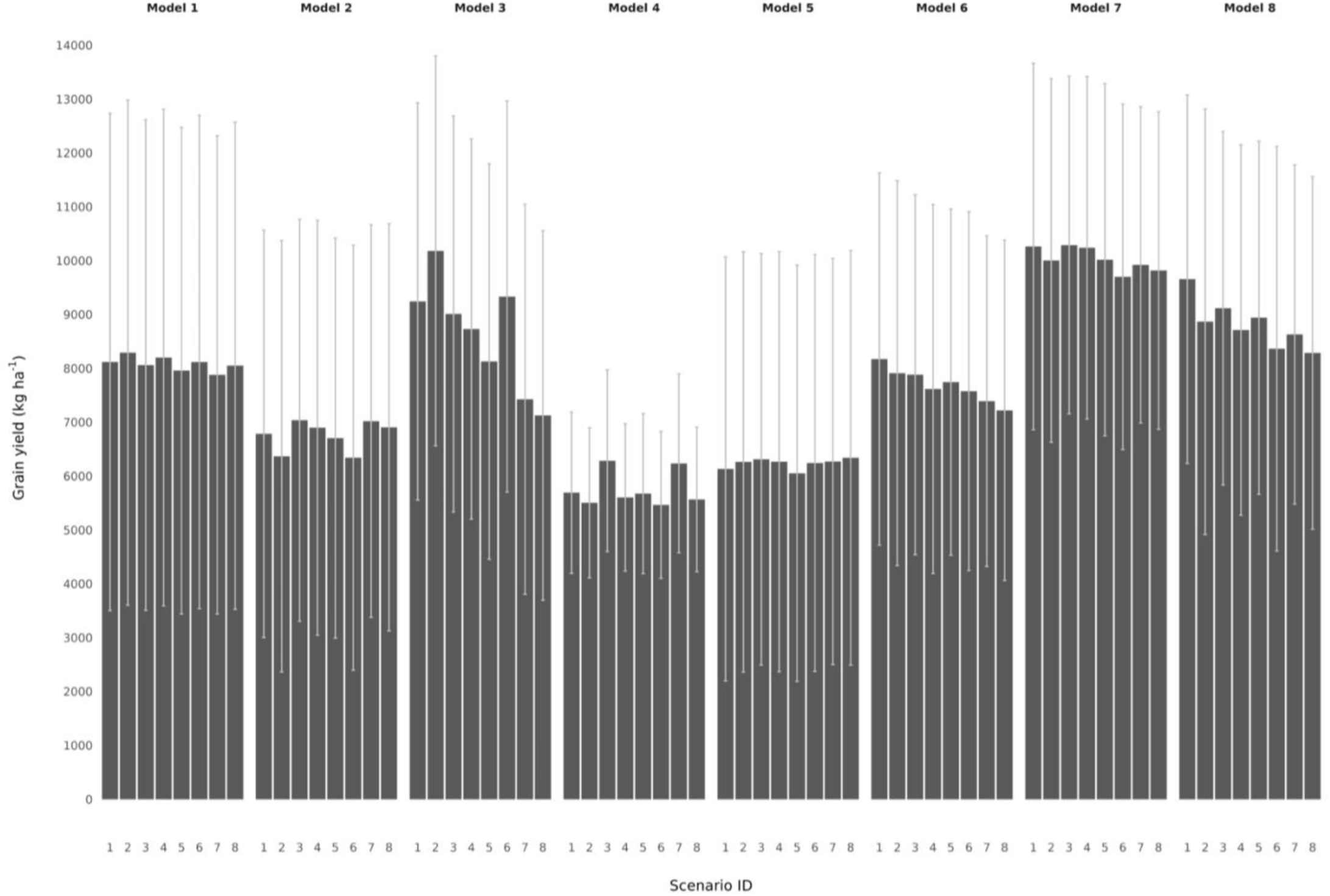
Mean maize grain yields and standard deviations across CSMs (from Model 1 to Model 8, at the top) and scenarios (from 1 to 8 at the bottom). The means are calculated across the entire Midwest region, encompassing 40,000 unique identifiers (UIDs), over a period of 42 years of simulation.

**Fig. S4.**
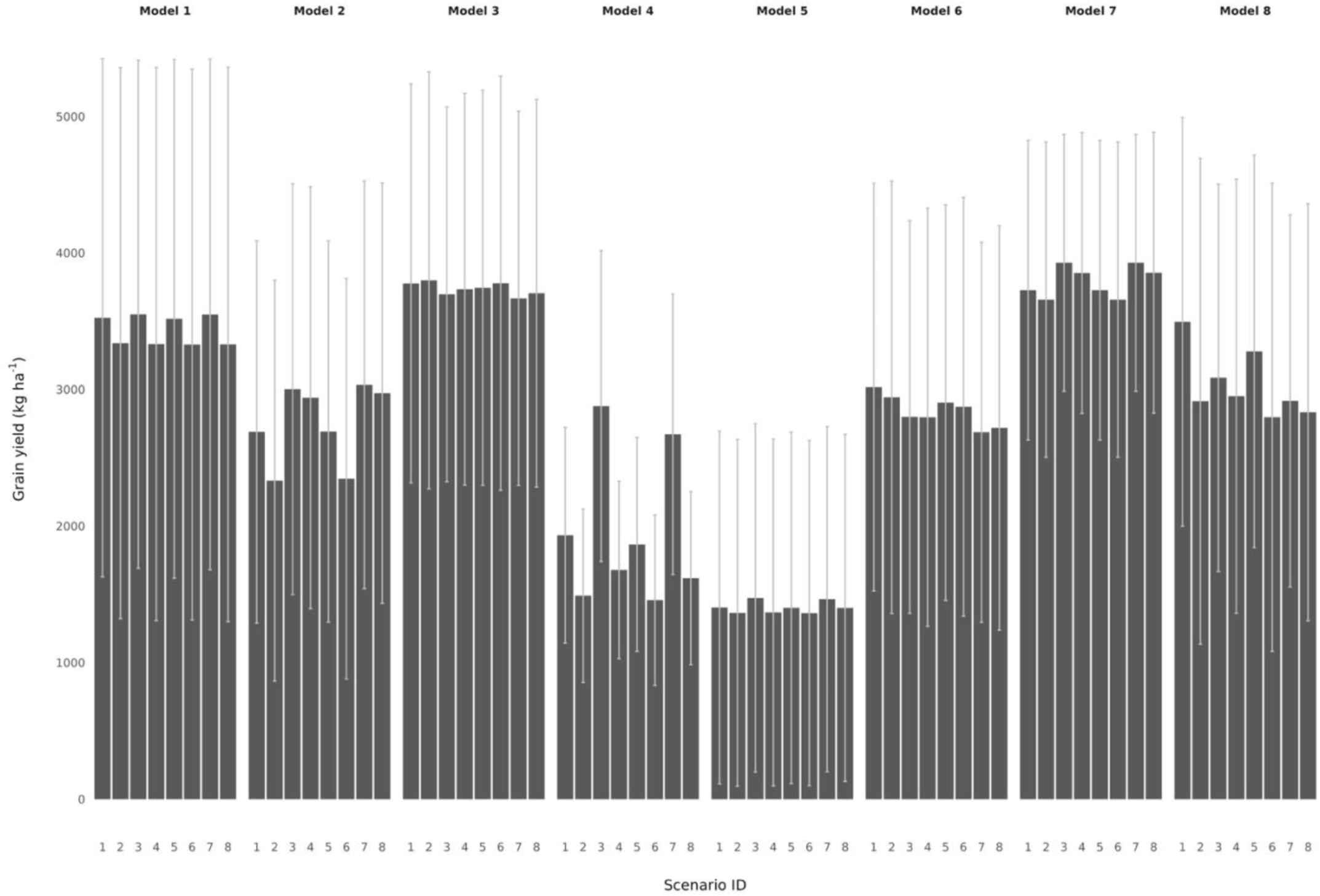
Mean soybean grain yields and standard deviations across CSMs (from Model 1 to Model 8, at the top) and scenarios (from 1 to 8 at the bottom). The means are calculated across the entire Midwest region, encompassing 40,000 unique identifiers (UIDs), over a period of 42 years of simulation.

**Fig. S5.**
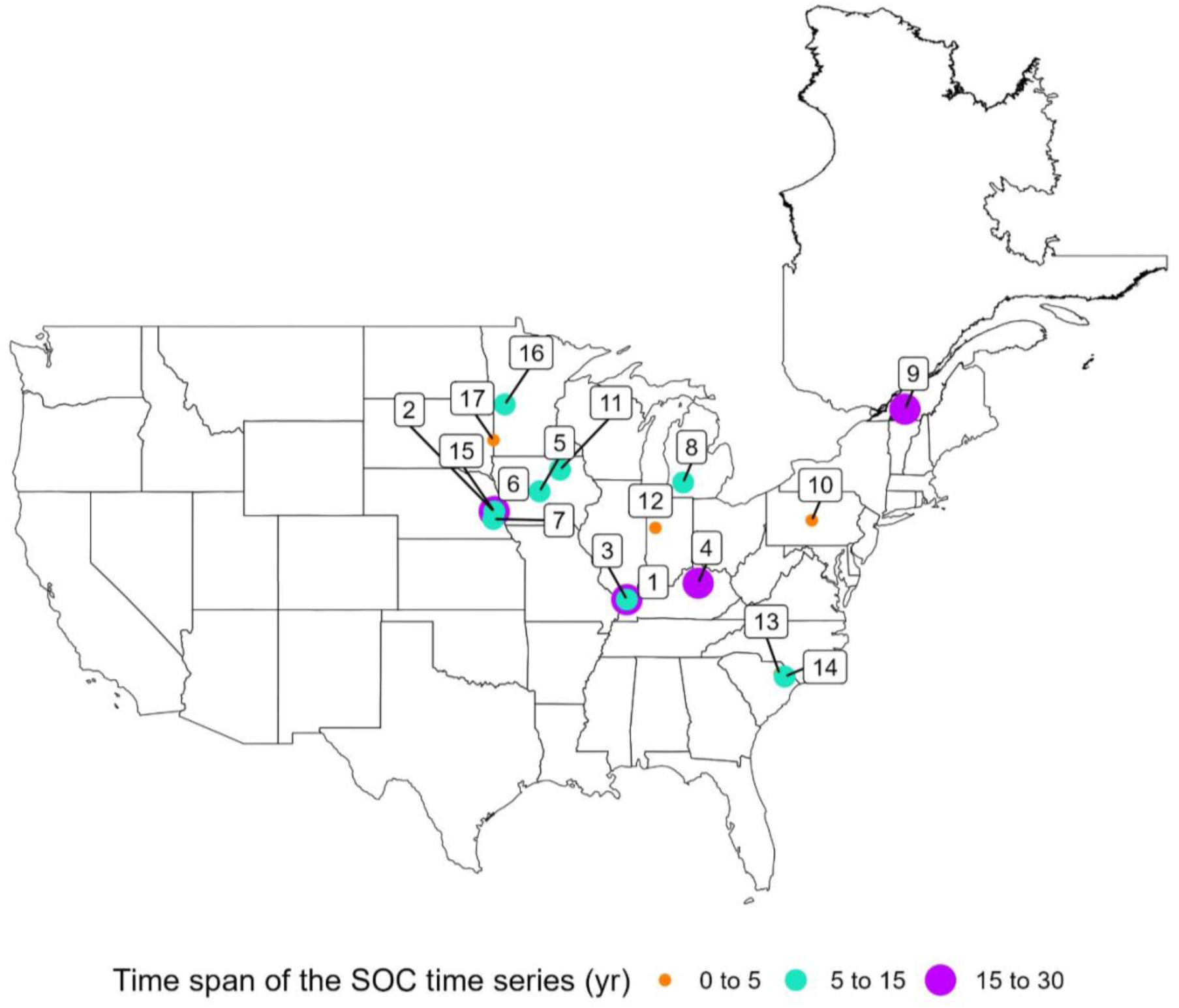
Map of the 17 experimental sites used for the Multi-model Ensemble validation. Time span of each experimental site is computed between the first and the last SOC samplings.

**Fig. S6.**
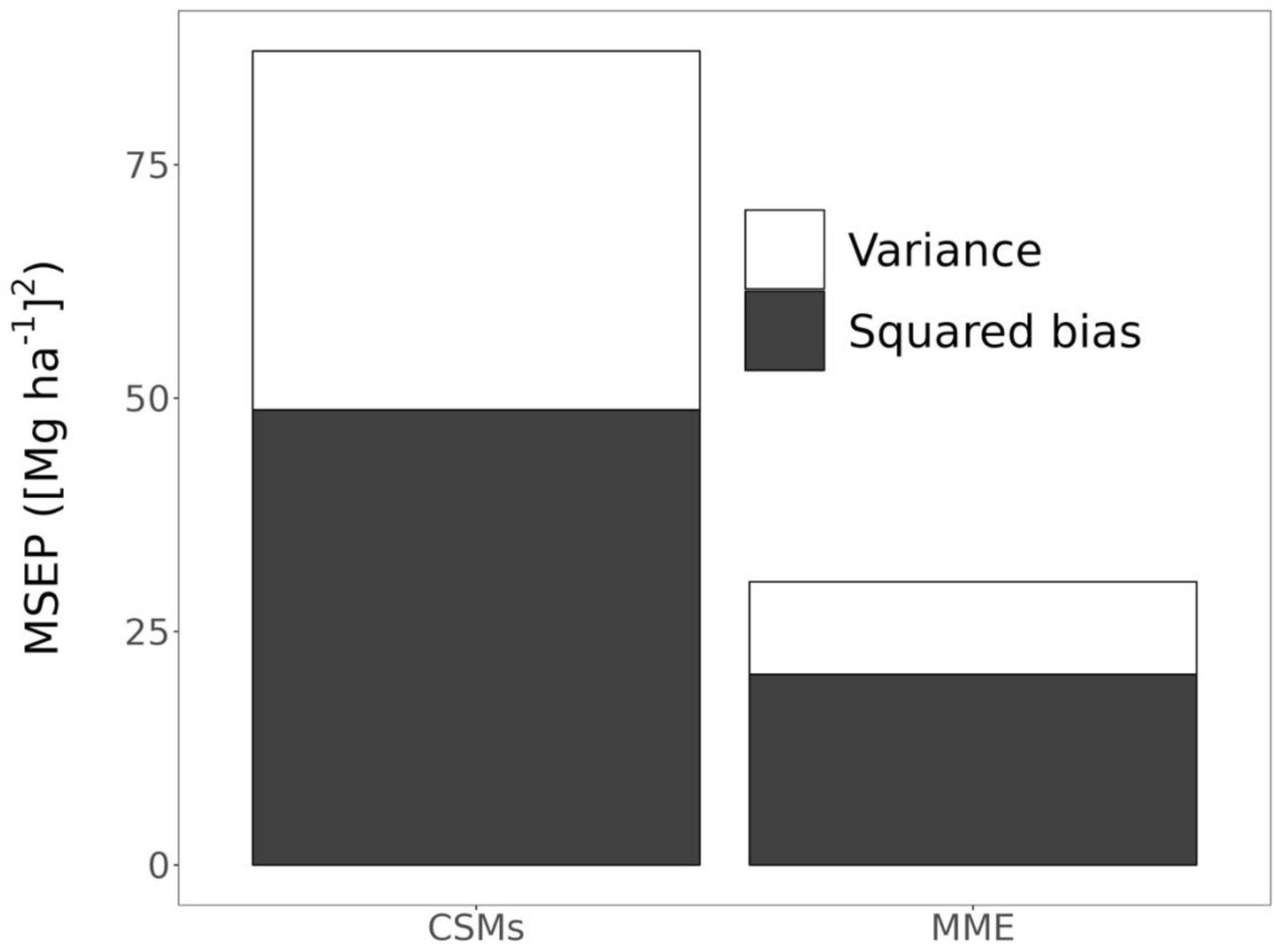
Uncertainty analysis conducted for the CSMs population and MME across the measured experimental sites and treatments. Uncertainty is expressed as the mean squared error of prediction (MSEP) and its decomposition in prediction squared bias and prediction variance on the raw SOC stock data, as reported in equations 6-8 under the material and methods paragraph. Lower uncertainty corresponds to lower MSEP values and vice versa.

**Fig. S7.**
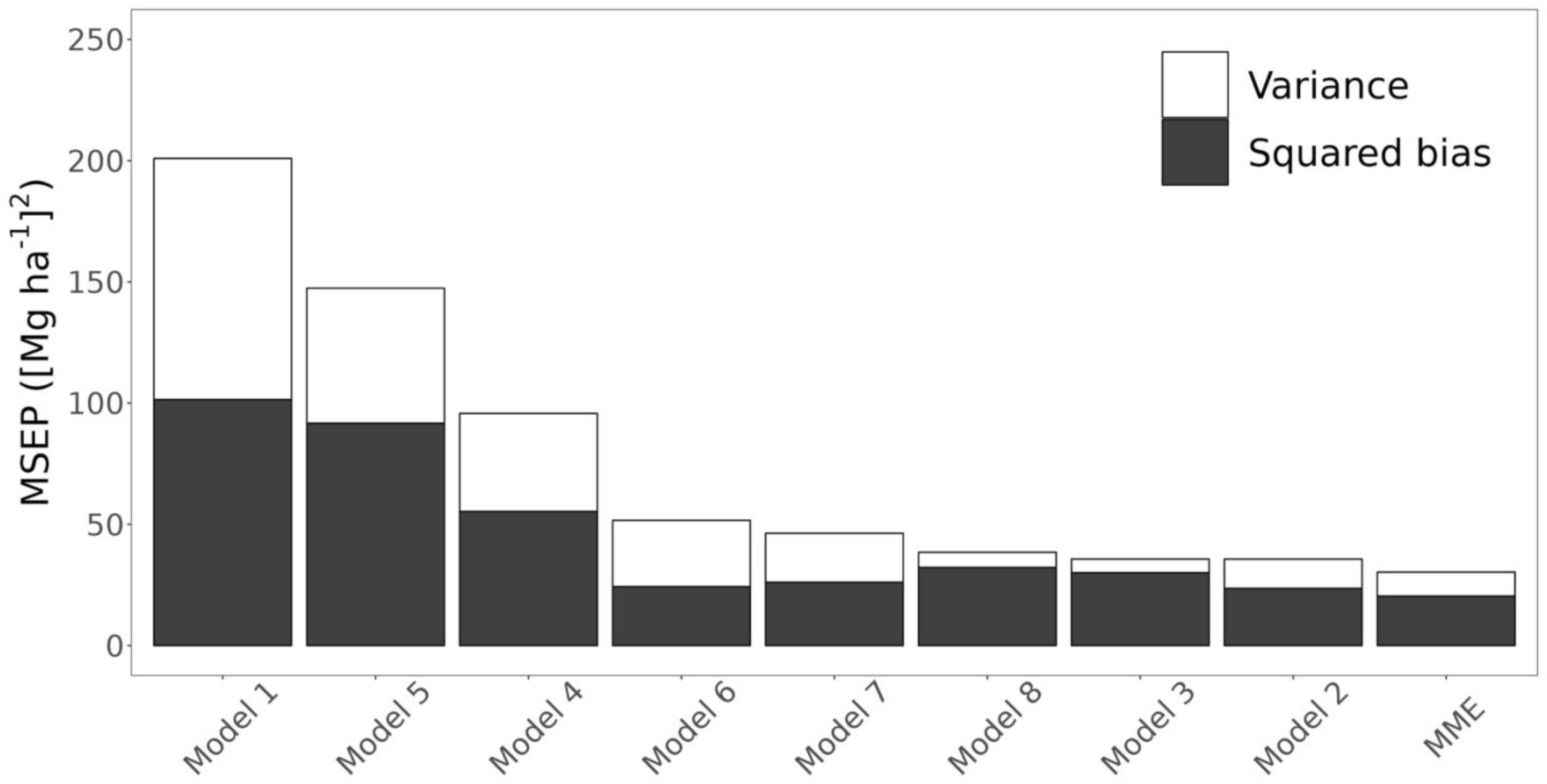
Uncertainty analysis conducted for each individual CSM and MME across the measured experimental sites and treatments. Uncertainty is expressed as the mean squared error of prediction (MSEP) and its decomposition in prediction squared bias and prediction variance on the raw SOC stock data, as reported in equations 6-8 under the material and methods paragraph. Lower uncertainty corresponds to lower MSEP values and vice versa.

**Fig. S8.**
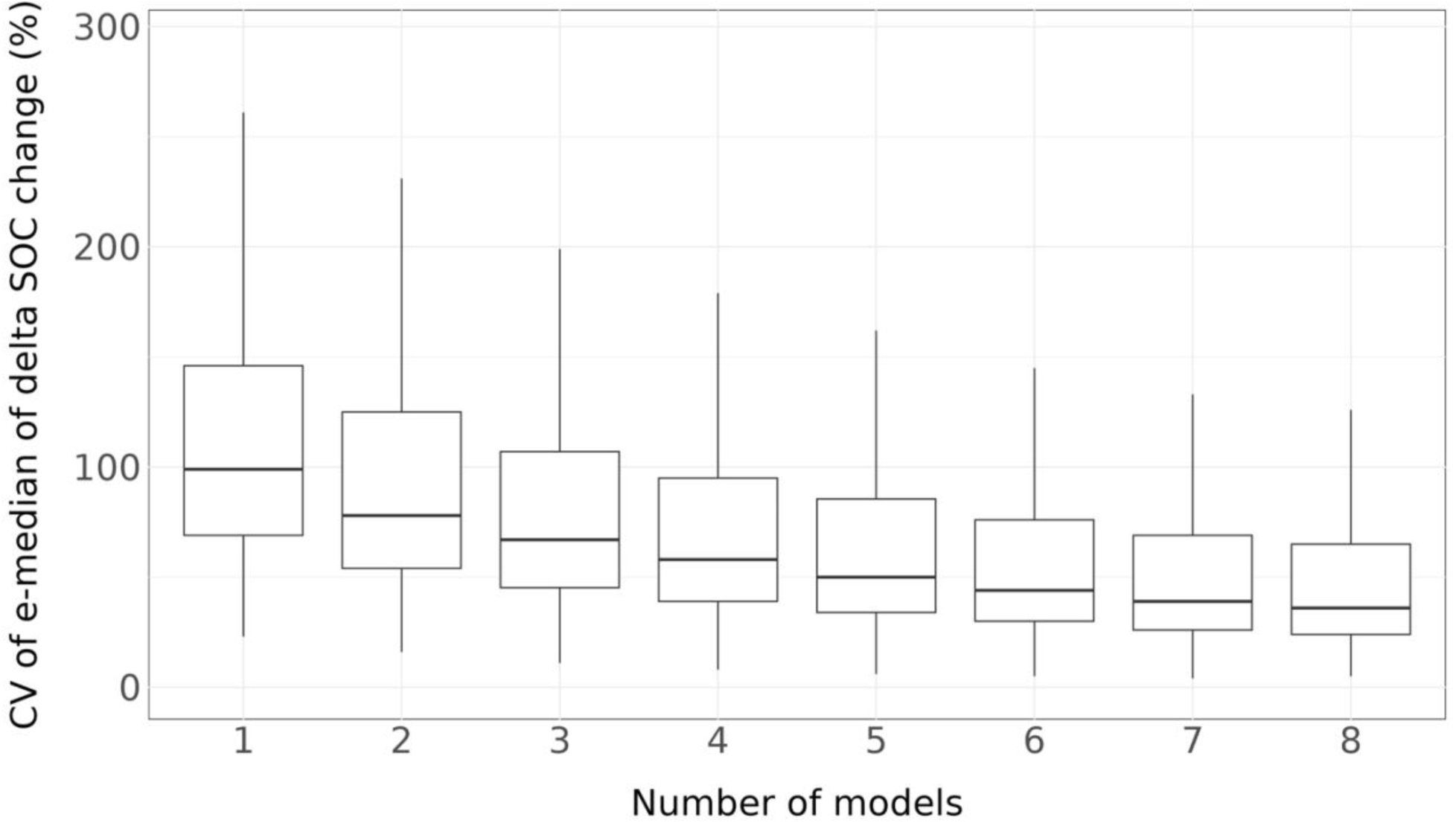
Boxplots of the coefficient of variation (CV) of the delta SOC change e-median across varying number of crop simulation models (CSM) involved in the multi-model ensemble (MME) computation over the US Midwest upscale. The final MME included all 8 models. Coefficient of variation values are retrieved from a bootstrap computation as reported in equation 9 in the material and method paragraph.

**Fig. S9.**
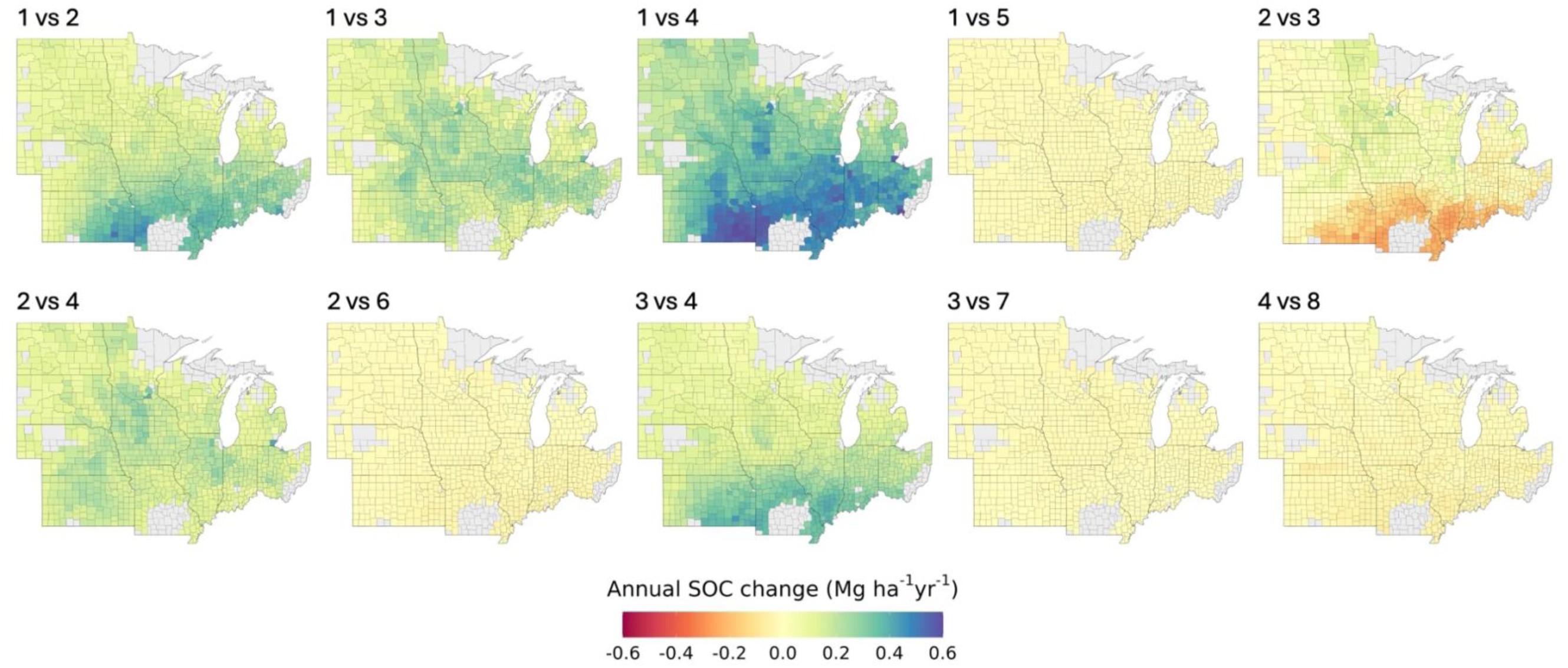
MME county mean annual delta SOC change (Mg ha^-1^ yr^-1^) in the 0-30 cm soil layer as a difference between different scenarios. Numbers on the top of each figure refer to the scenarios under comparison, with numbers from 1 to 8 indicating the scenario ID as reported in Table 1 (scenarios 1-4) and table S3 (scenarios 5-8).

**Fig. S10.**
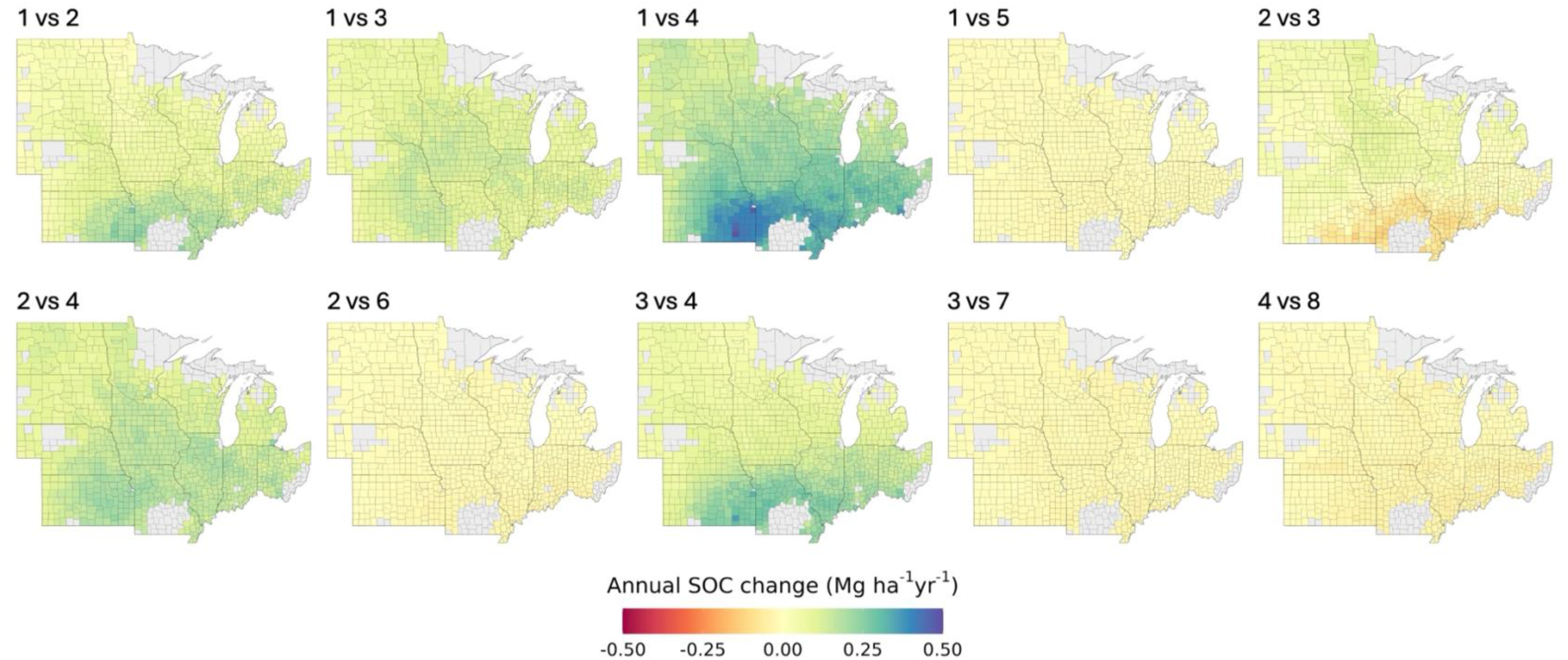
MME county mean annual delta SOC change (Mg ha^-1^ yr^-1^) in the 0-15 cm soil layer as a difference between different scenarios. Numbers on the top of each figure refer to the scenarios under comparison, with numbers from 1 to 8 indicating the scenario ID as reported in Table 1 (scenarios 1-4) and table S3 (scenarios 5-8).

**Fig. S11.**
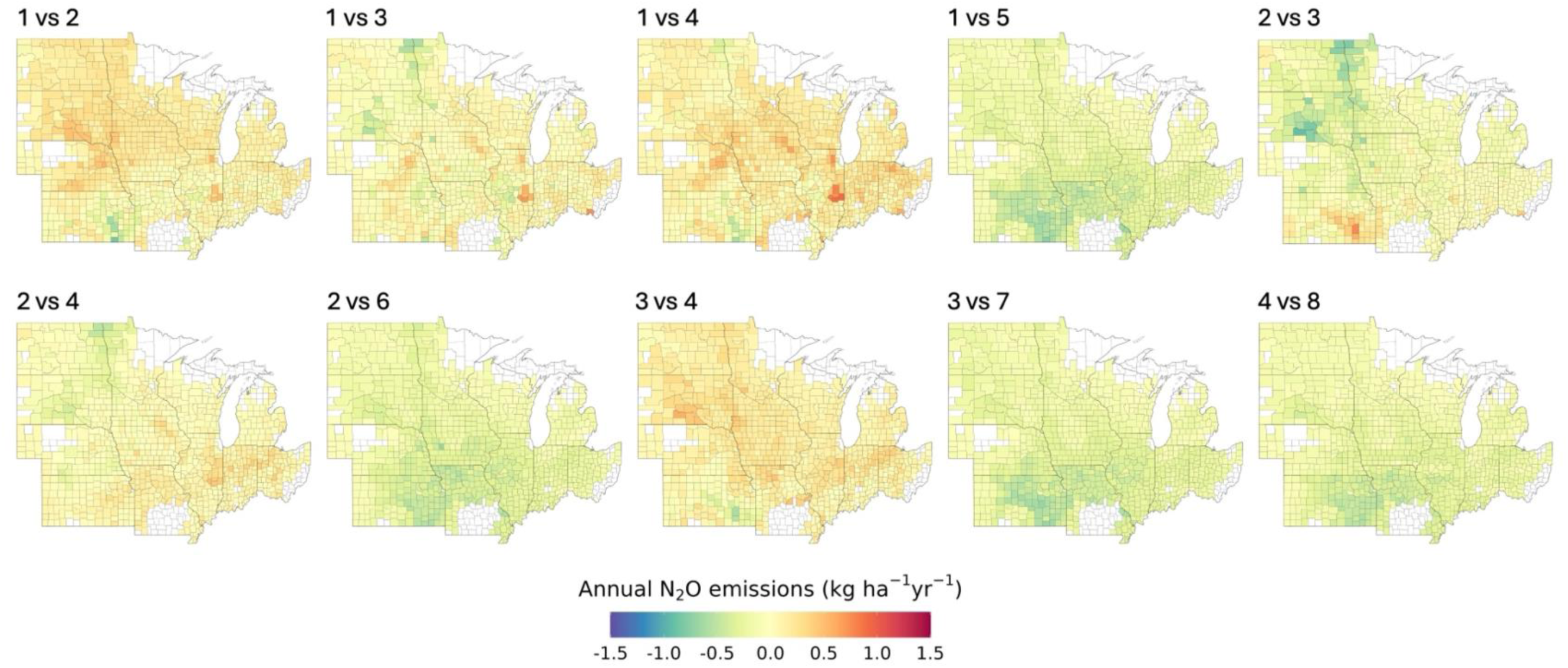
MME county mean annual N2O emission (kg ha^-1^ yr^-1^) as a difference between different scenarios. Numbers on the top of each figure refer to the scenarios under comparison, with numbers from 1 to 8 indicating the scenario ID as reported in Table 1 (scenarios 1-4) and table S3 (scenarios 5-8).

**Fig. S12.**
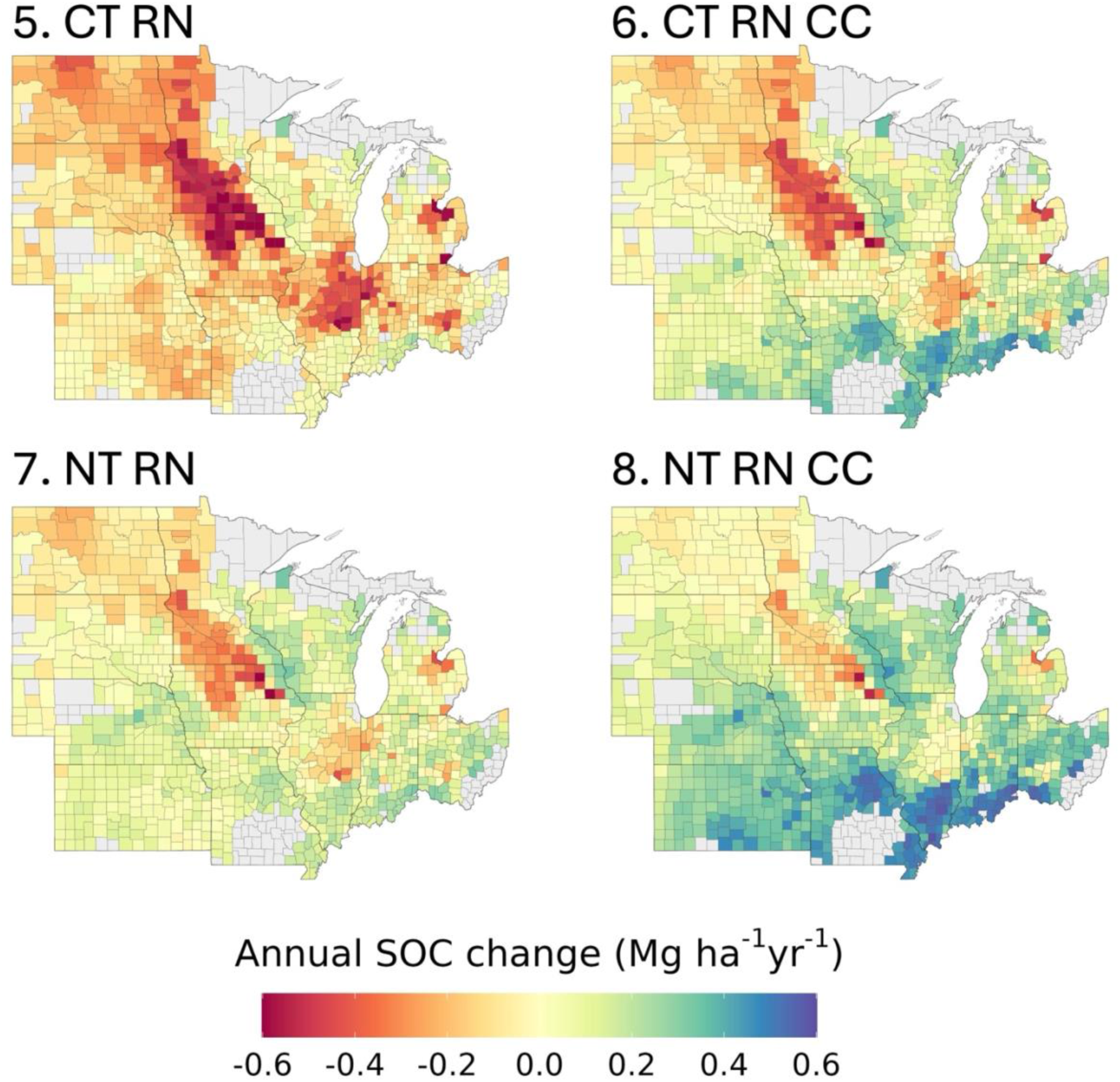
MME county mean annual delta SOC change (Mg ha^-1^ yr^-1^) in the 0-30 cm soil layer across the 12 states considered in the analysis. Each map represents a single scenario, with numbers from 5 to 8 indicating the scenario ID as reported in table S3. Scenarios from 1 to 4 are reported in the main paper.

**Fig. S13.**
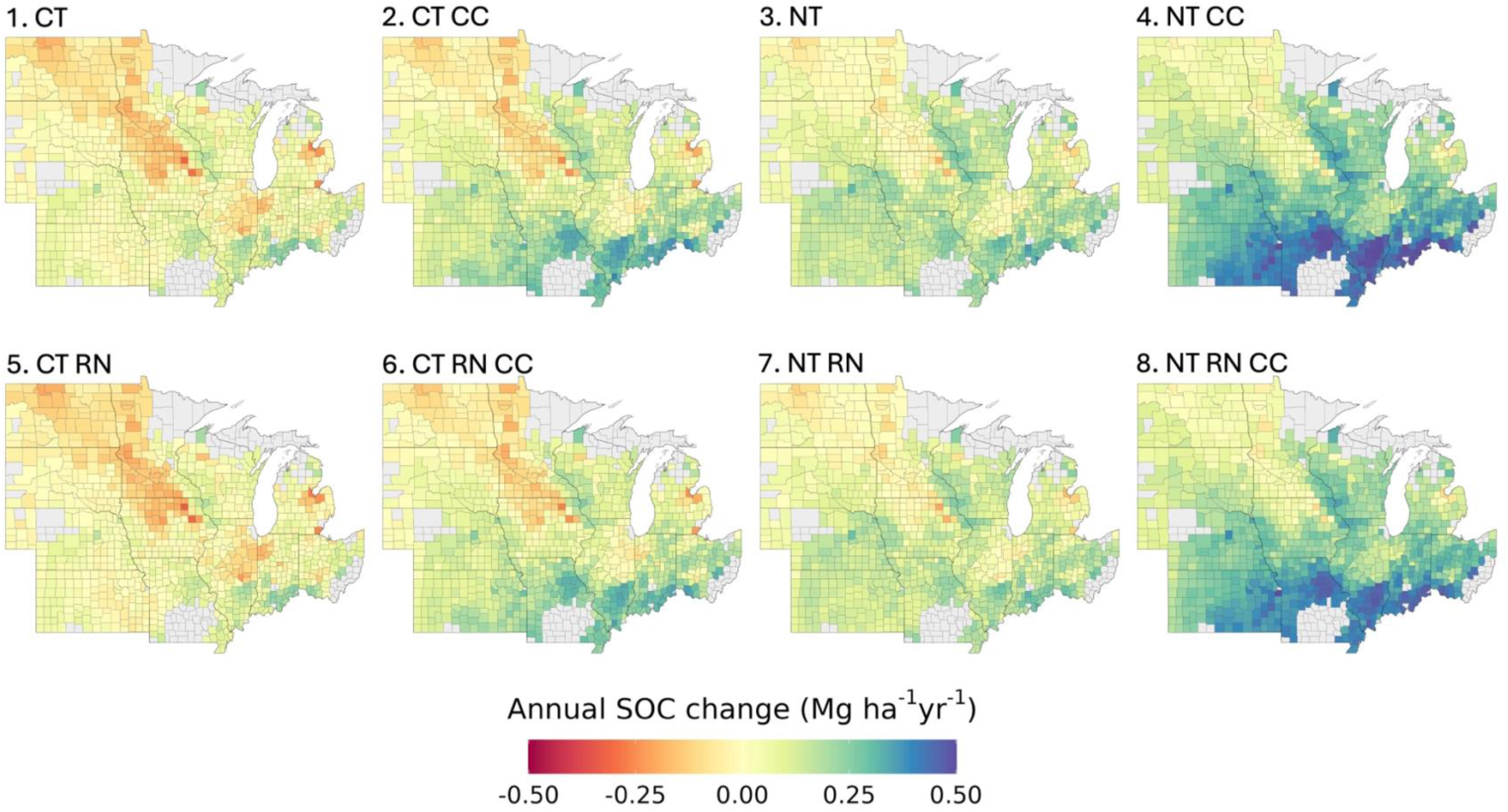
MME county mean annual delta SOC change (Mg ha^-1^ yr^-1^) in the 0-15 cm soil layer across the 12 states considered in the analysis. Each map represents a single scenario, with numbers from 1 to 8 indicating the scenario ID as reported in Table 1 (scenarios 1-4) and table S3 (scenarios 5-8).

**Fig. S14.**
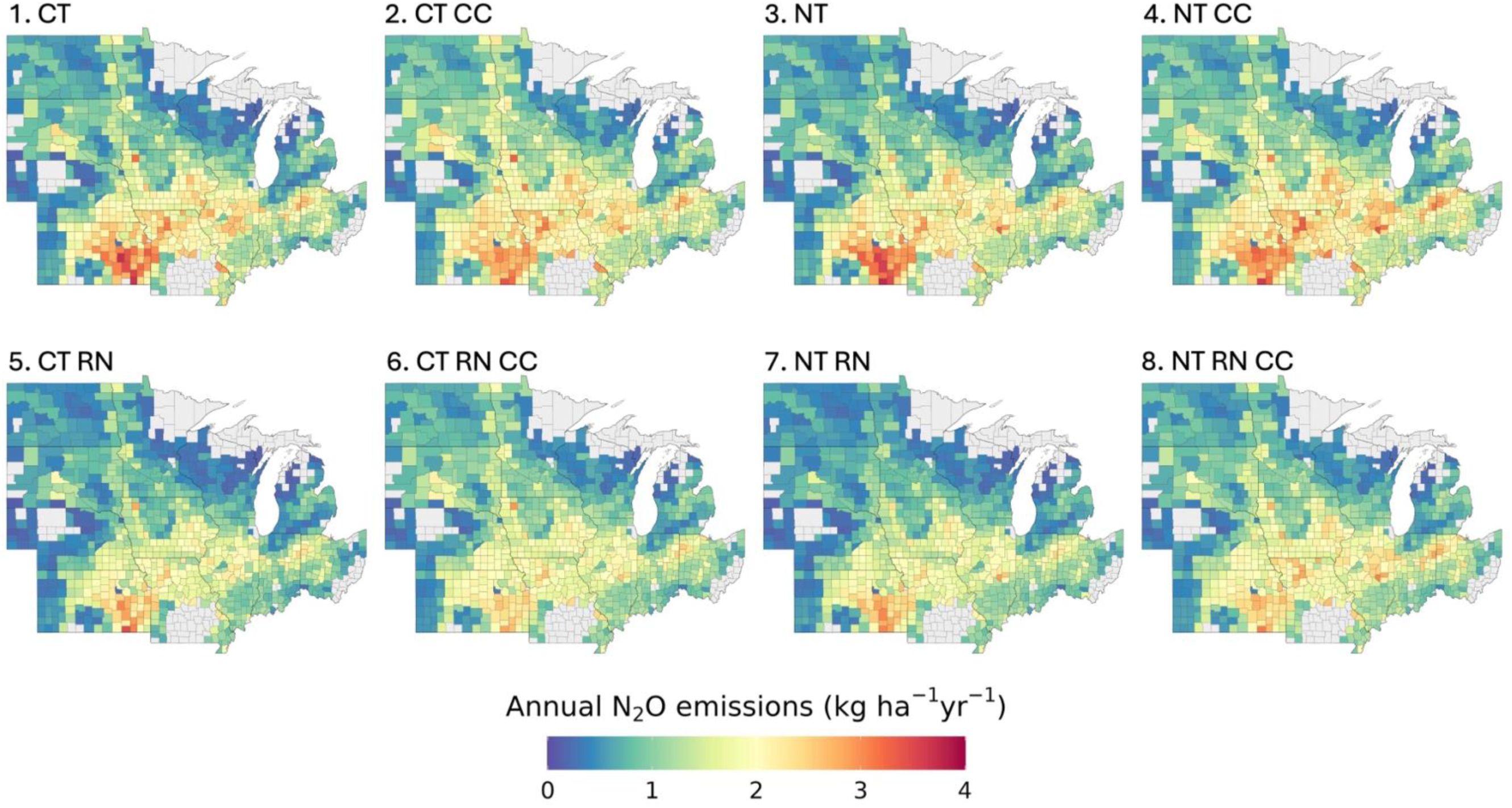
MME county mean cumulative annual N2O emission (kg ha^-1^ yr^-1^) across the 12 states considered in the analysis. Each map represents a single scenario, with numbers from 1 to 8 indicating the scenario ID as reported in Table 1 (scenarios 1-4) and table S3 (scenarios 5- 8).

**Fig. S15.**
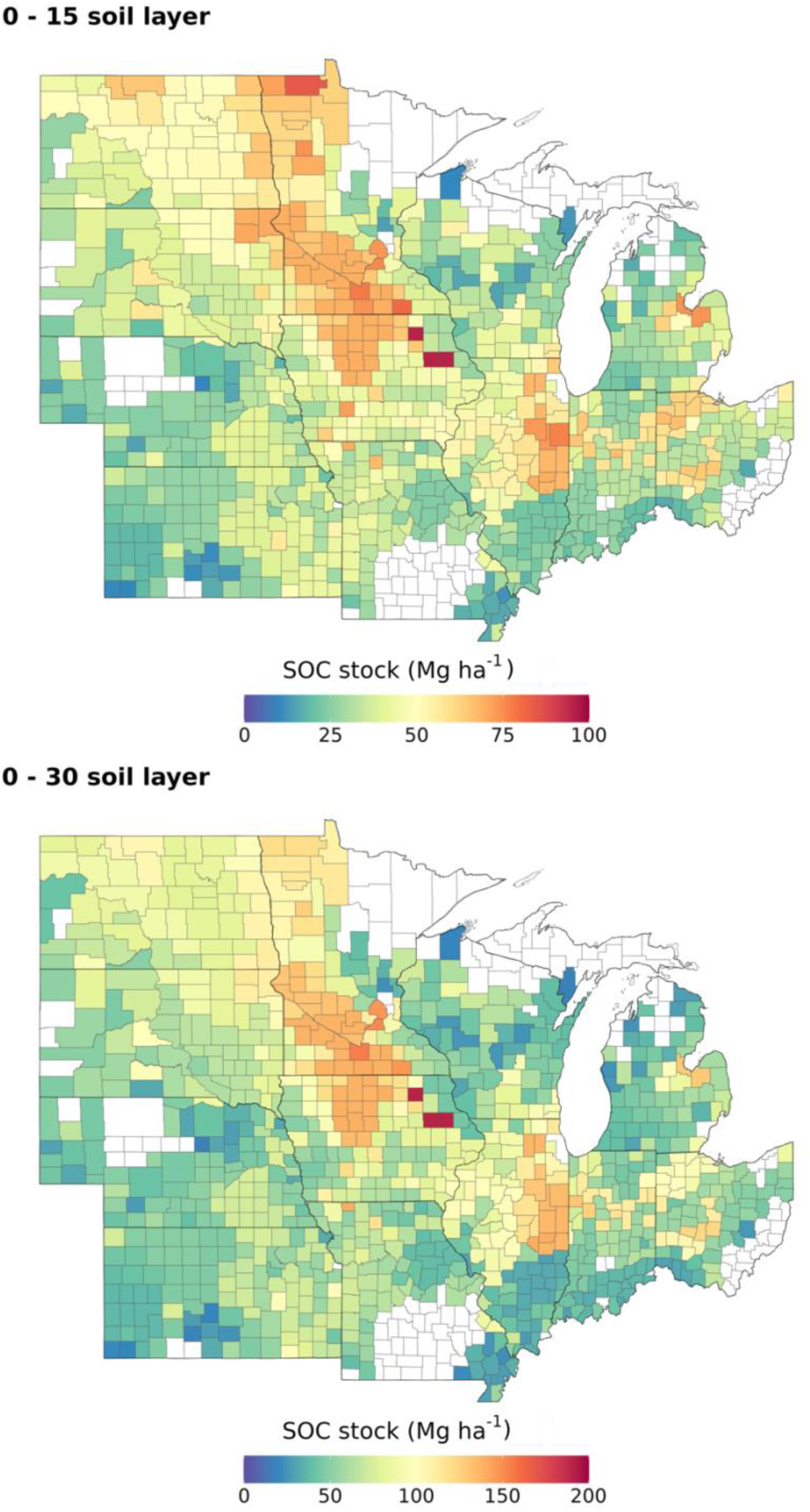
Initial SOC stock (Mg ha^-1^) at 0-15 cm (top) and 0-30 cm (bottom) cm soil layer expressed as a county mean.

**Fig. S16.**
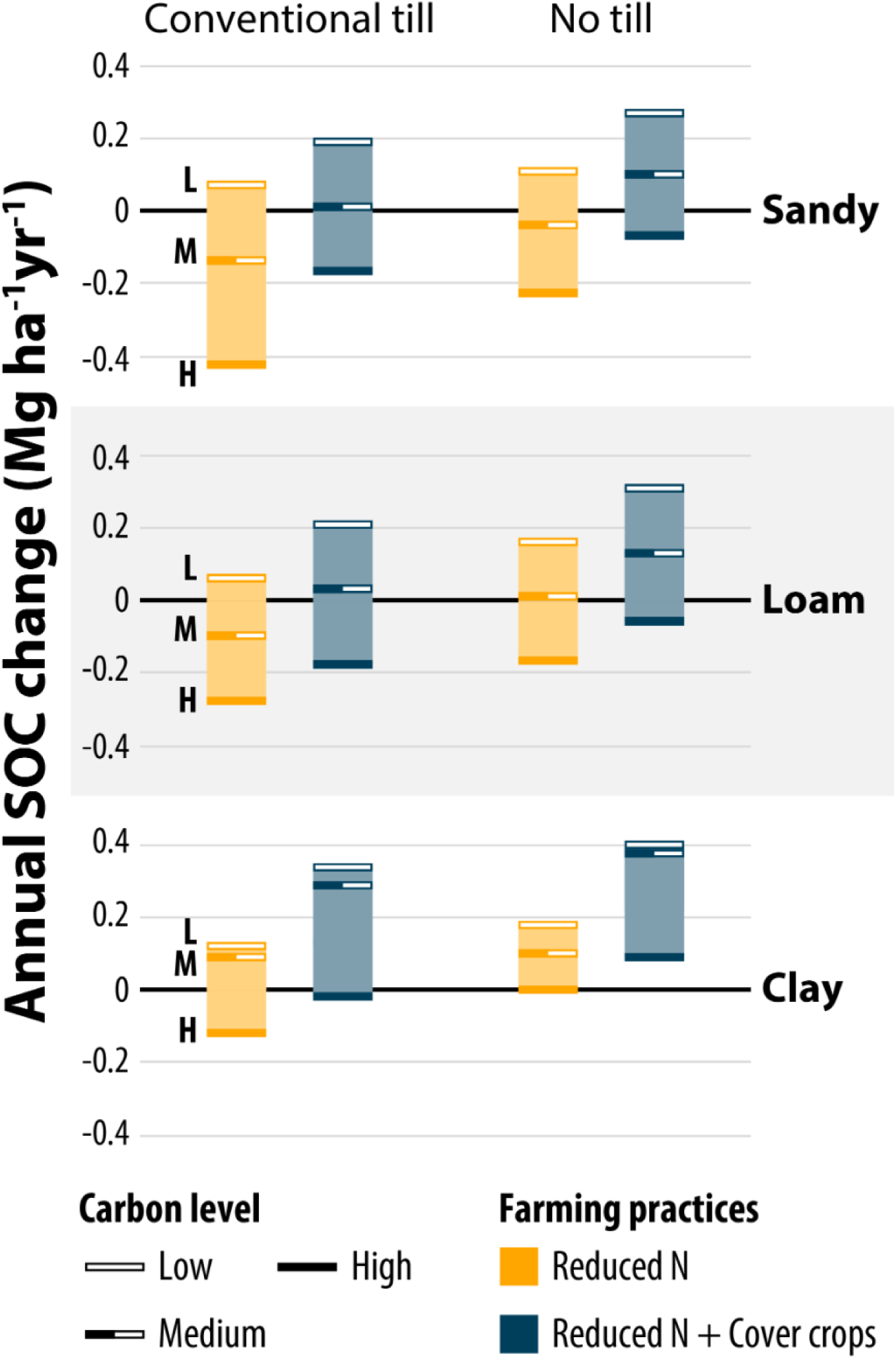
MME soil organic carbon (SOC) change (Mg ha^-1^ yr^-1^, 0-30 cm) across scenarios (5 to 8, from left to right), soil texture (Sandy, Loam and Clay), and initial SOC stock level (Low, < 40 Mg C ha^-1^; Medium, 40-80 Mg C ha^-1^; and High, > 80 Mg C ha^-1^). Annual SOC change values derived from all UIDs mean across the US Midwest. All scenarios use maize-soybean crop rotation.

**Fig. S17.**
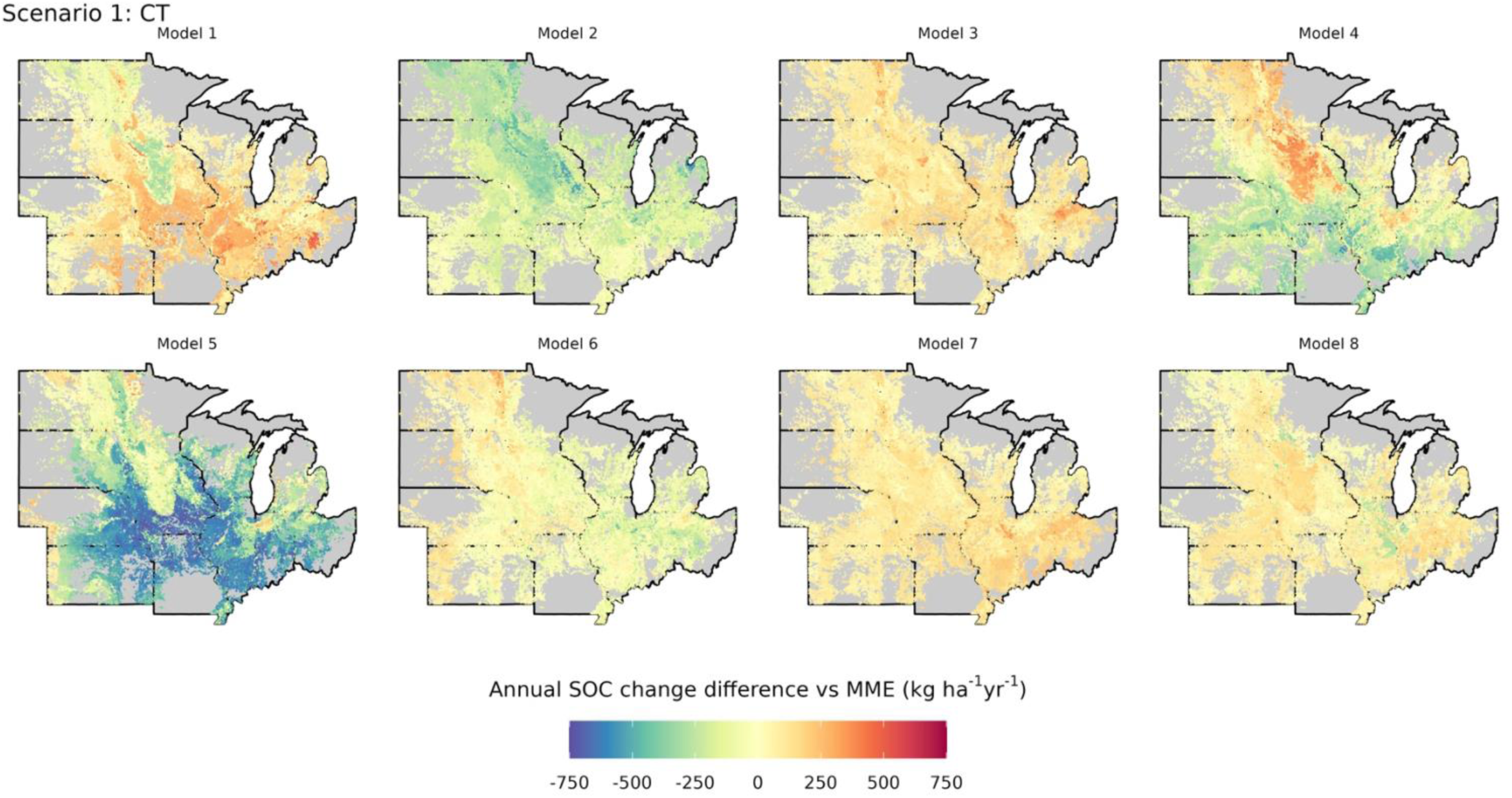
Annual SOC change difference between single CSM and MME in the 0-15 cm soil layer under Scenario 1 (CT) for each UID included in the study (n = 40,000). Each map represents a single CSM, with numbers from 1 to 8 indicating the model ID.

**Fig. S18.**
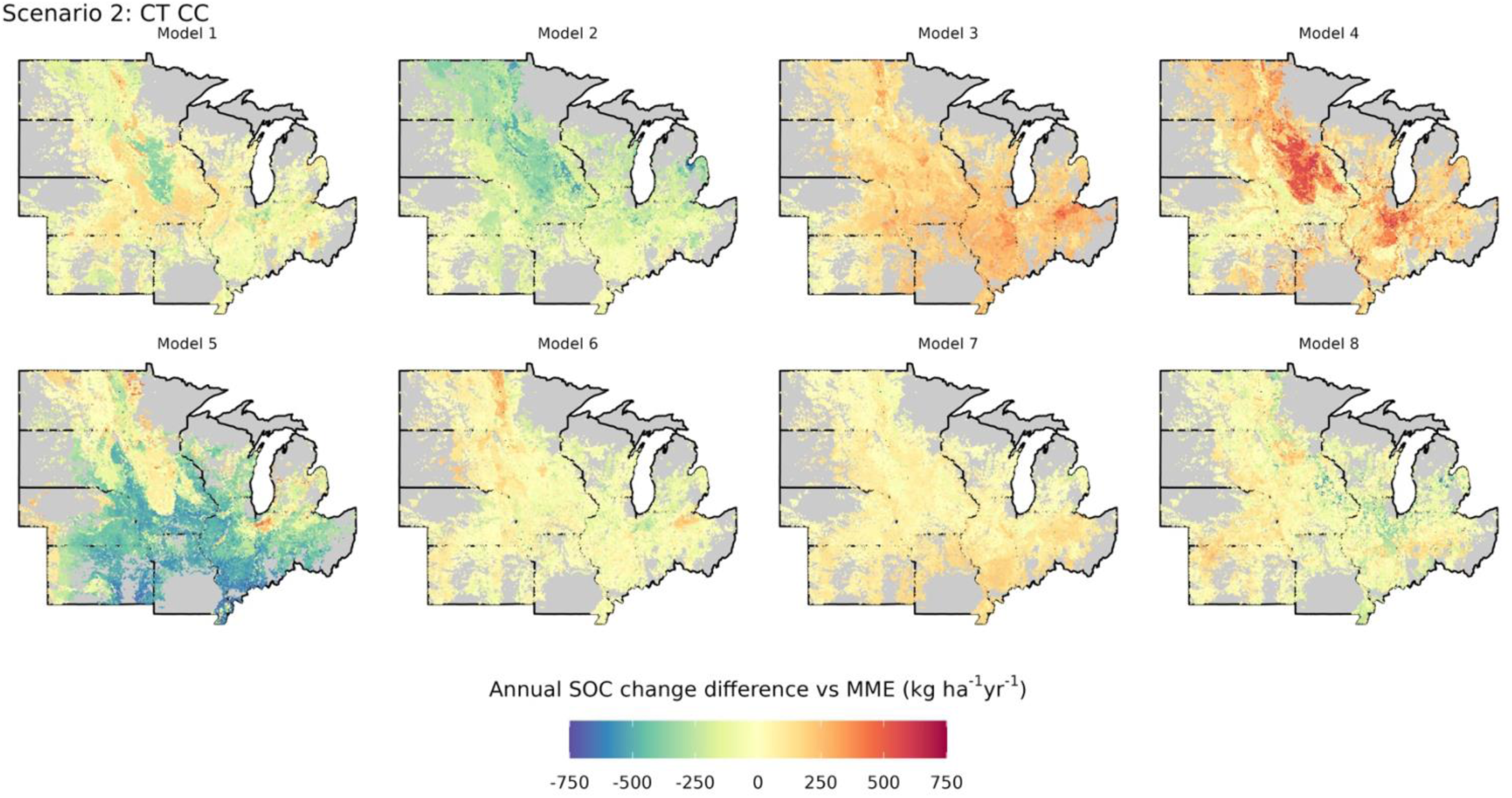
Annual SOC change difference between single CSM and MME in the 0-15 cm soil layer under Scenario 2 (CT CC) for each UID included in the study (n = 40,000). Each map represents a single CSM, with numbers from 1 to 8 indicating the model ID.

**Fig. S19.**
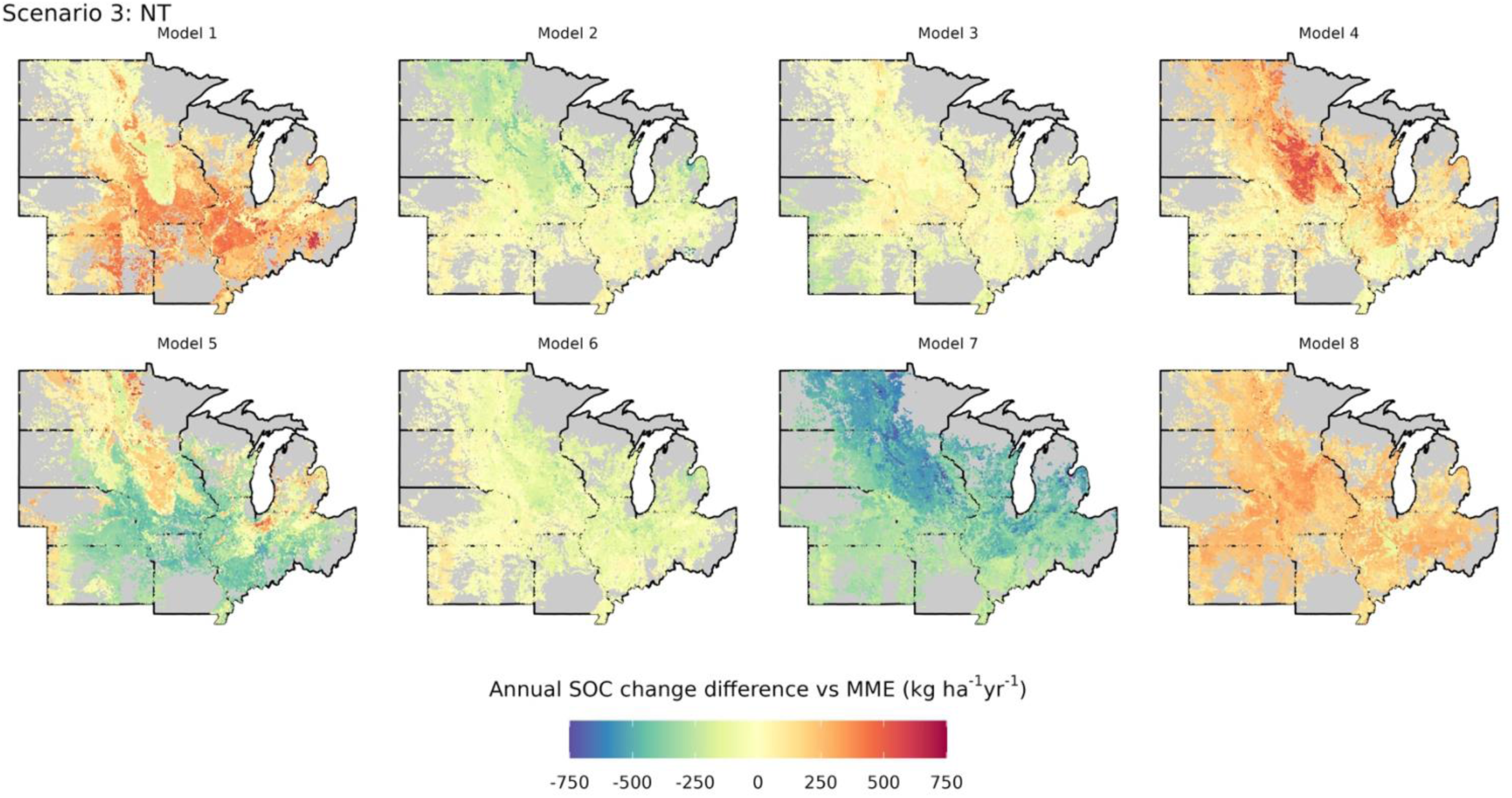
Annual SOC change difference between single CSM and MME in the 0-15 cm soil layer under Scenario 3 (NT) for each UID included in the study (n = 40,000). Each map represents a single CSM, with numbers from 1 to 8 indicating the model ID.

**Fig. S20.**
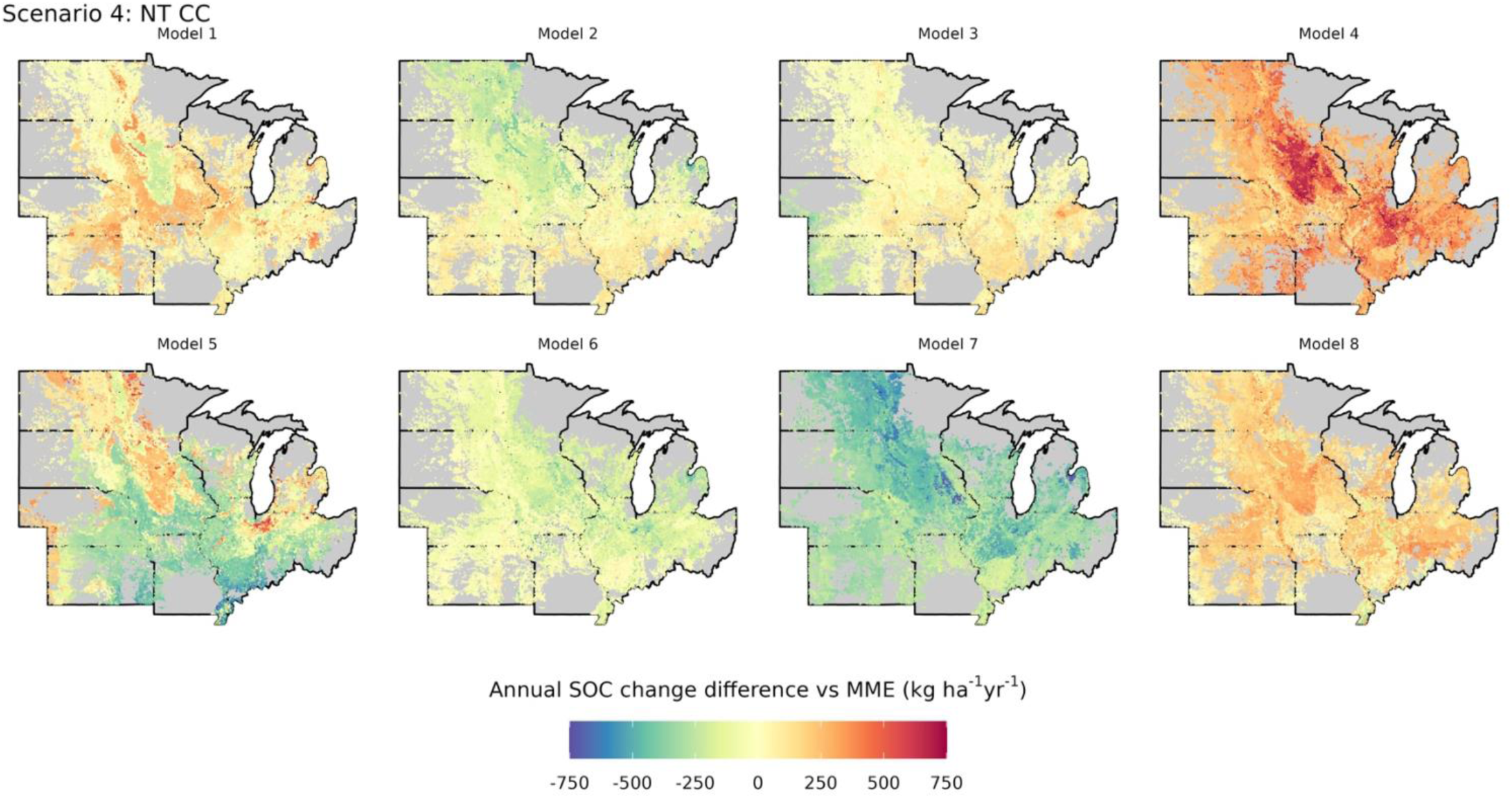
Annual SOC change difference between single CSM and MME in the 0-15 cm soil layer under Scenario 4 (NT CC) for each UID included in the study (n = 40,000). Each map represents a single CSM, with numbers from 1 to 8 indicating the model ID.

**Fig. S21.**
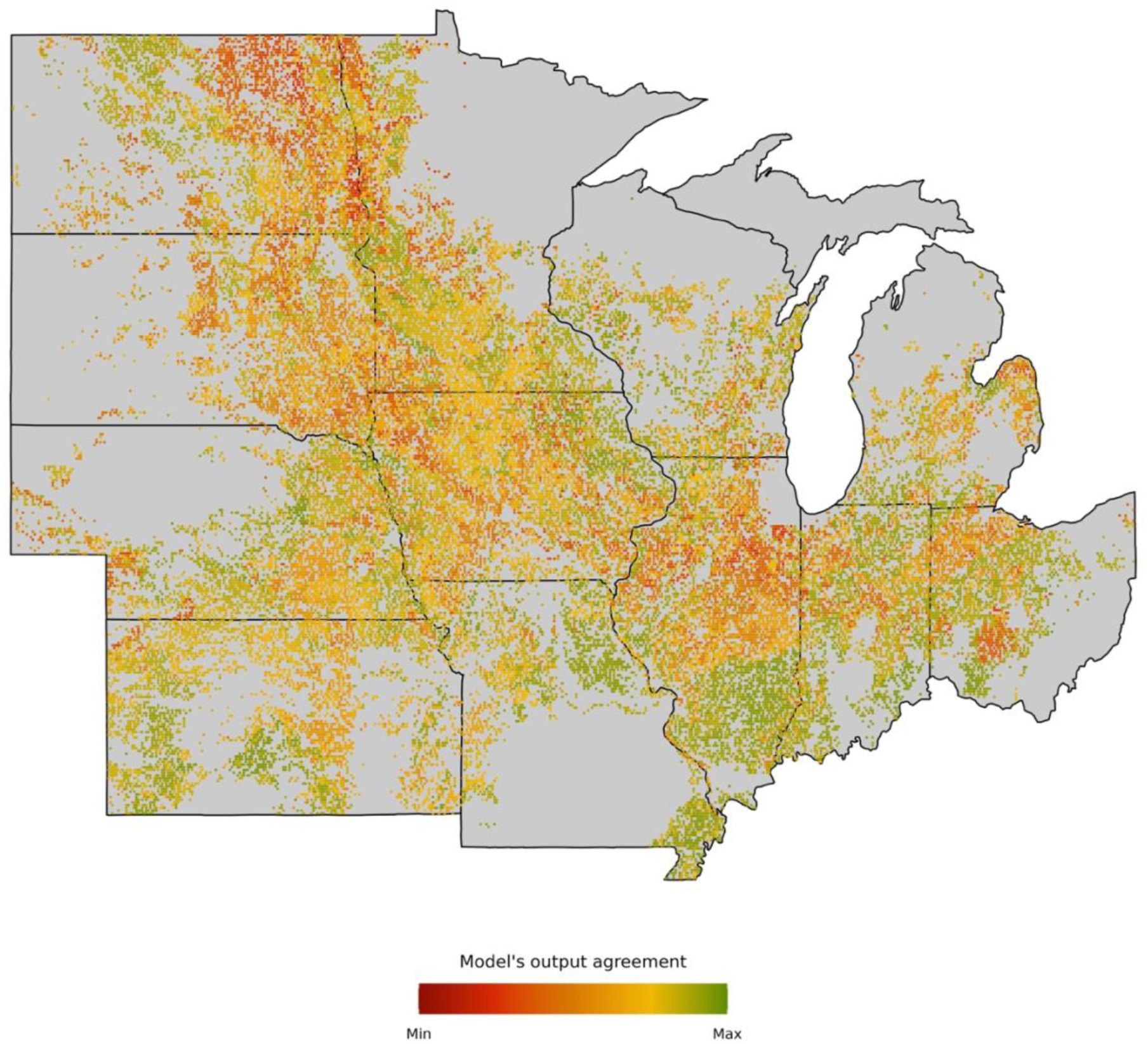
Overall CSMs output agreement across the 40000 locations using the delta SOC change as a target variable. Data comes from a normalized RRMSE (between 0 and 1) computed following equation 3 (see Material and Methods) where predicted values represent MME and observed data are represented by CSMs output. Normalized RRMSE values were averaged across scenarios (1 to 4) to obtain a single value per UID. Maximum agreement values (towards green) represent lower RRMSE and minimum agreement values (towards red) represent higher RRMSE.

### 3. SUPPLEMENTAL TABLES

**Table S1.** Process-based models used in the study, along with their main references and links to the official documentation.

| **Model** | **Version** | **Main reference** | **URL for documentation** |
| --- | --- | --- | --- |
| APSIM | 7.10 | Keating et al., 2003 | https://www.apsim.info/documentation/model-documentation/ |
| ARMOSA | rv1473 | Perego et al., 2013 | Available upon request |
| CROPSYST | 4 | Stöckle et al., 2003 | http://bioearth.wsu.edu/cropsyst_model.html |
| DAYCENT | vDev2016 | Parton et al., 1998 | https://www.nrel.colostate.edu/projects/daycent/ |
| DSSAT | 4.8 | Jones et al., 2003 | www.DSSAT.net |
| EPIC | 11.02 | Izaurrealde et al., 2006 | https://epicapex.tamu.edu/manuals-and-publications/ |
| SALUS | 3 | Basso and Ritchie, 2015 | https://basso.ees.msu.edu/salus/index.html |
| STICS | V10 | Beaudoin et al., 2023 | https://stics.inrae.fr/ |

**Table S2.**
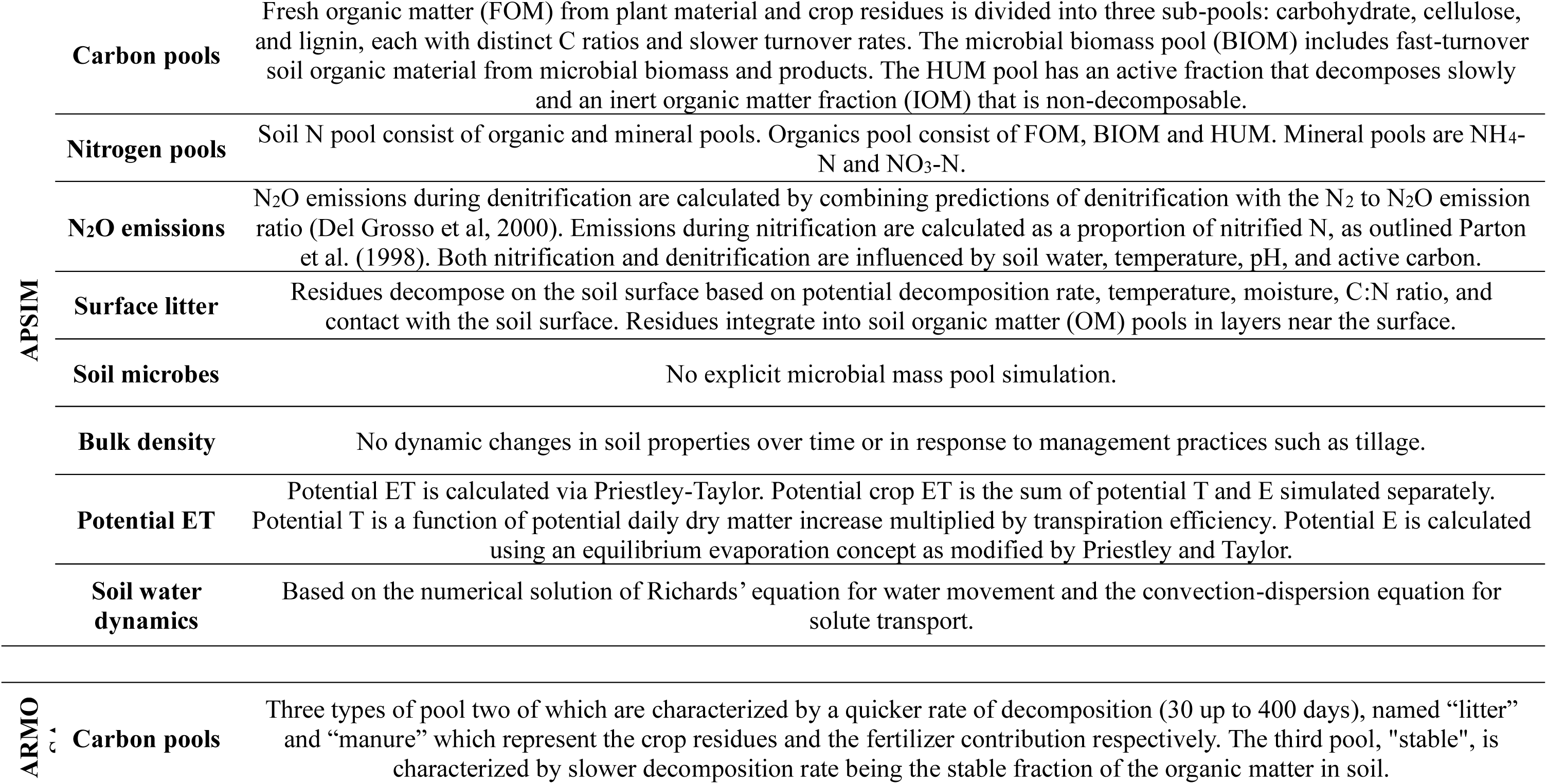
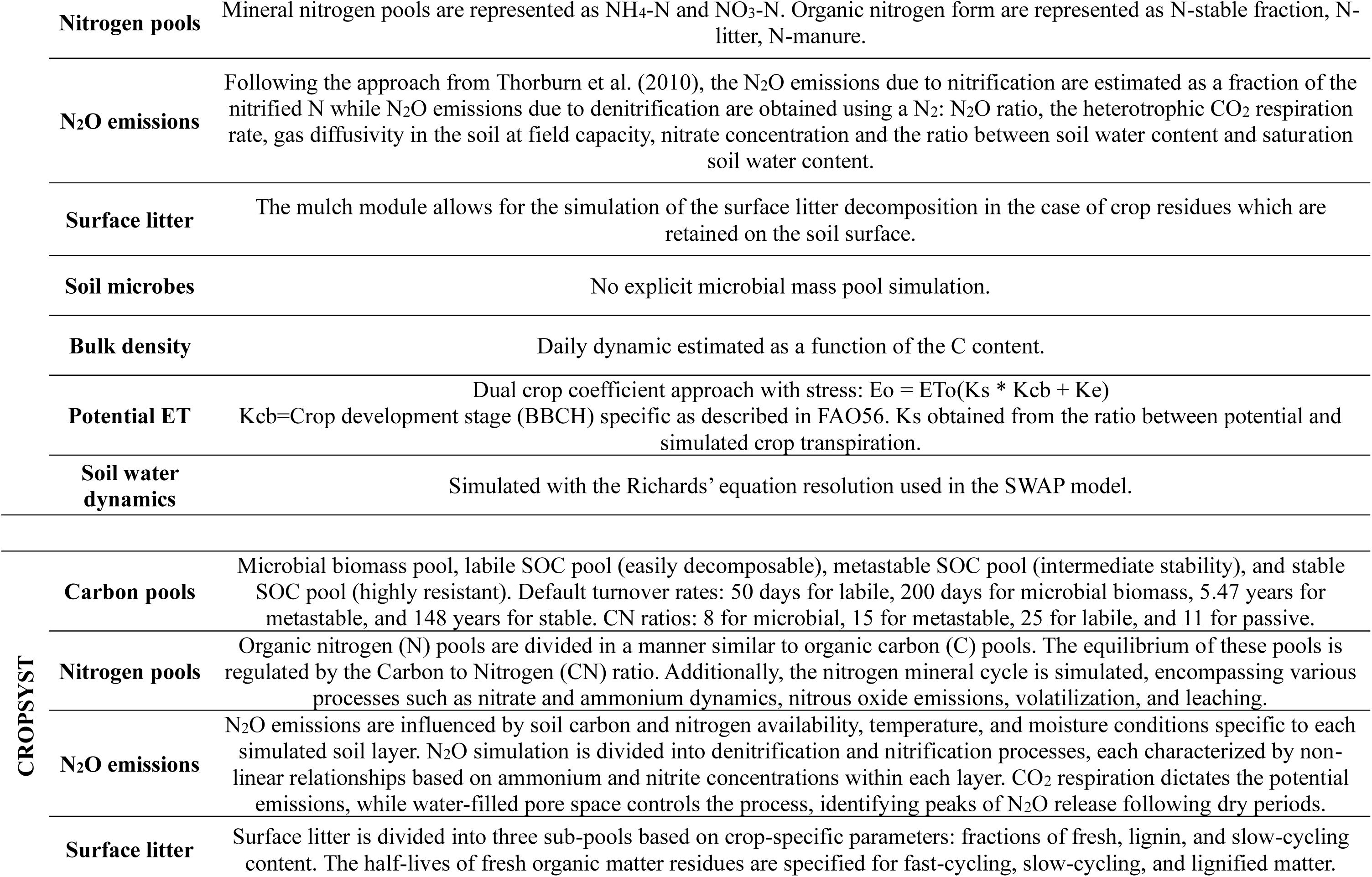
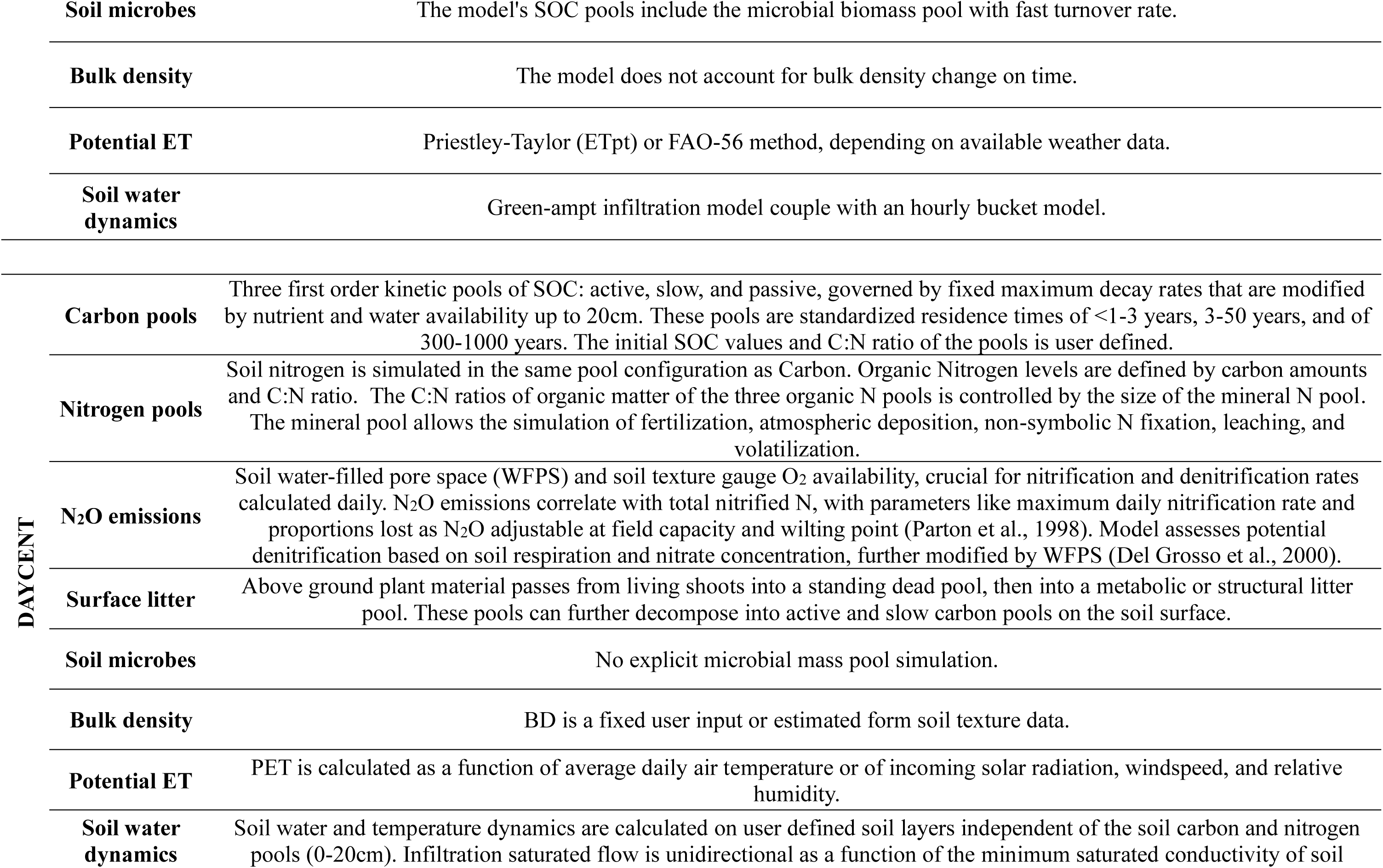
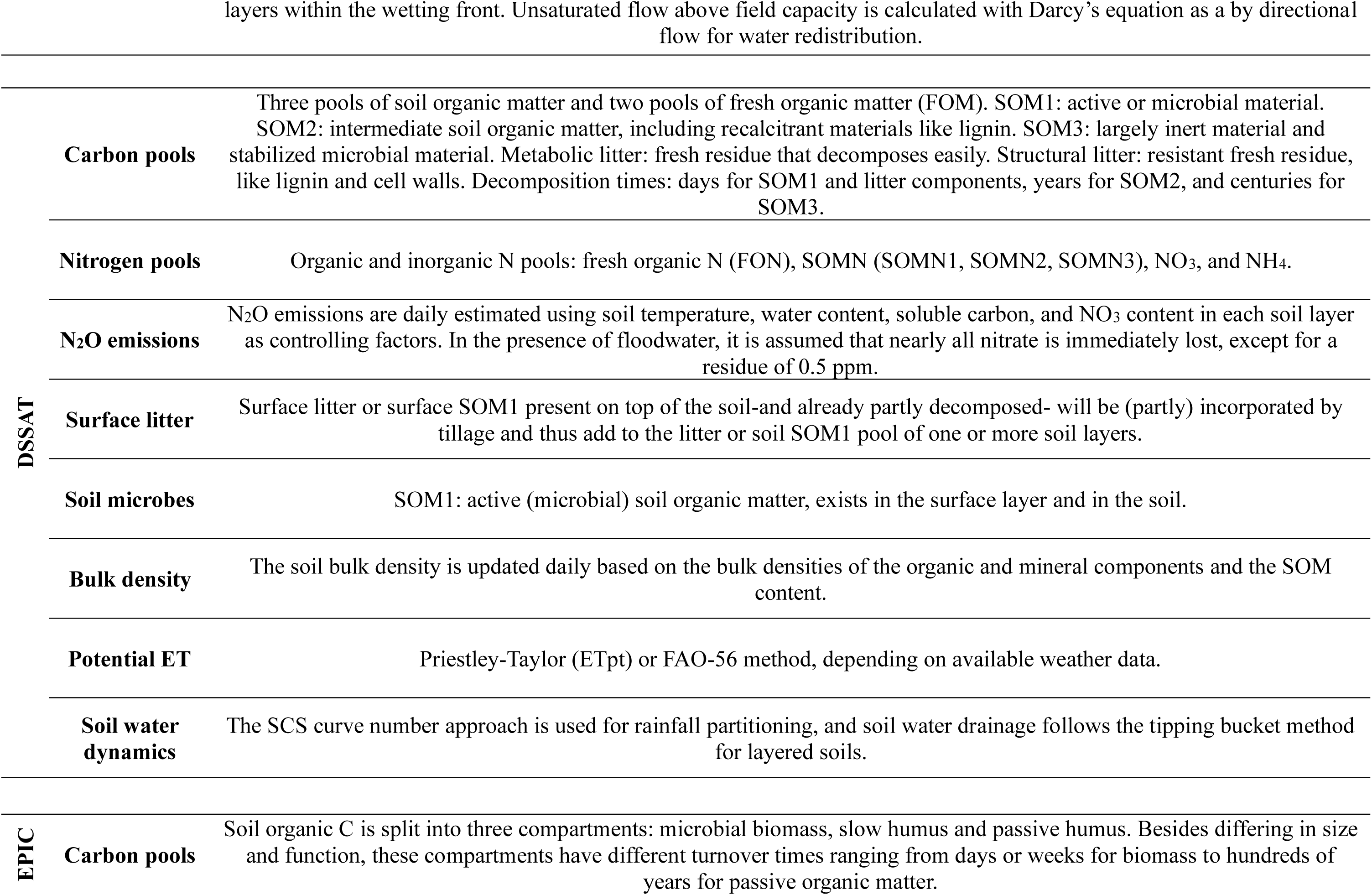
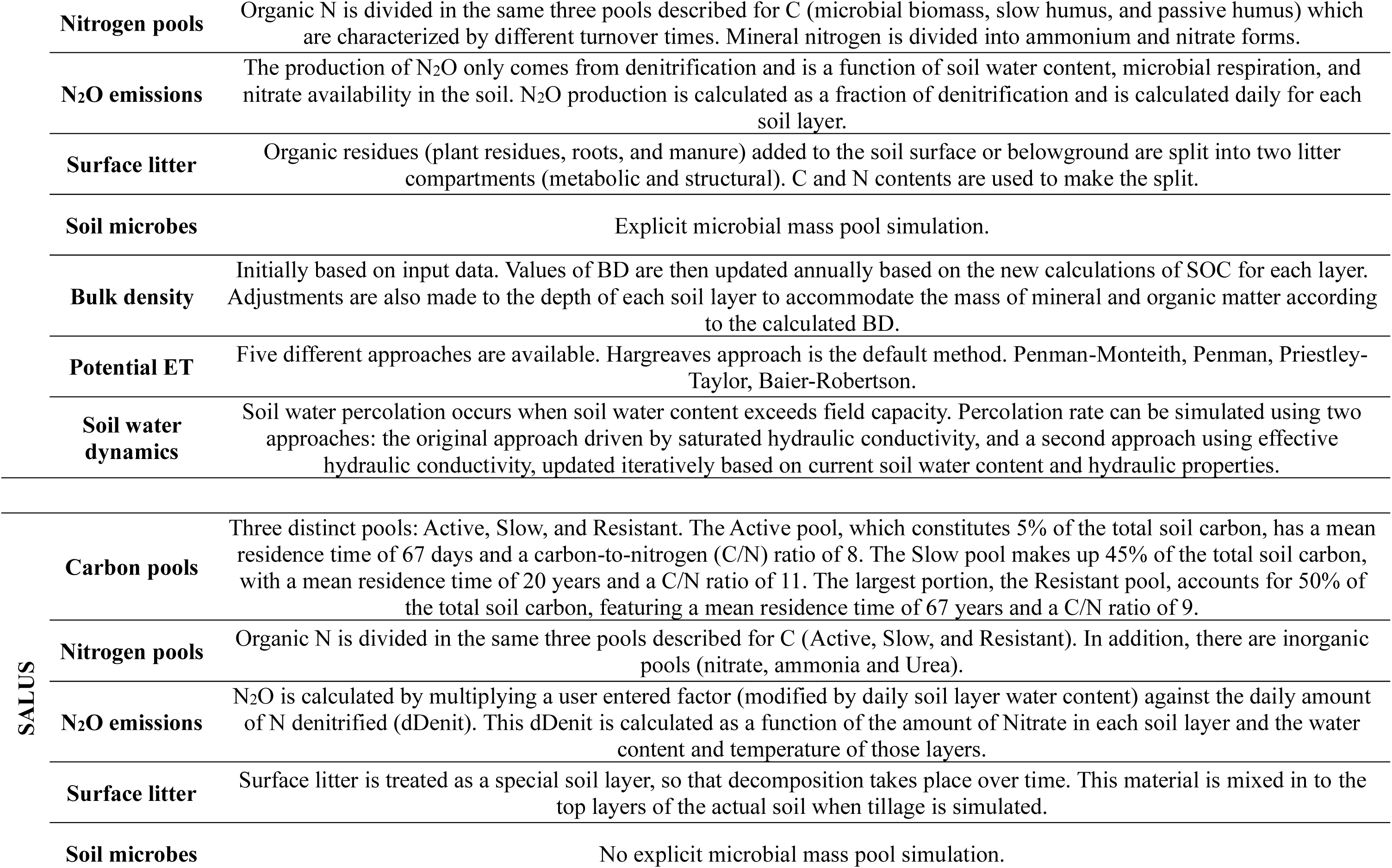
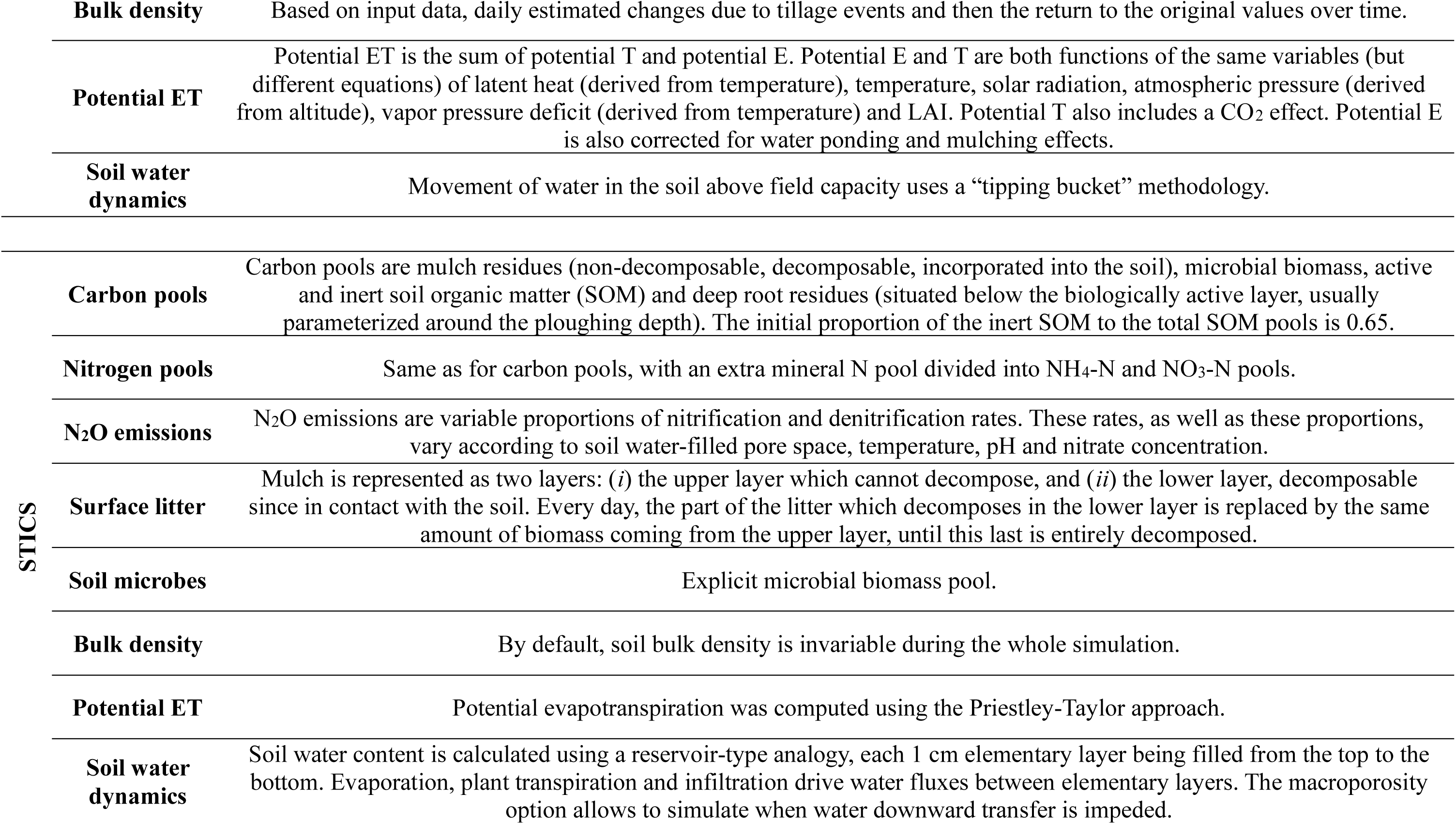
Summary of the models’ relevant features to this study. The table details each model with respect to: ’Carbon Pools’ (number and characteristics), ’Nitrogen Pools’ (number and characteristics), ’N2O emissions’ (description of N2O computation), ’Surface Litter’ (description of its decomposition process), ’Soil Microbes’ (presence of a microbe pool), ’Bulk Density’ (dynamic or fixed evolution), ’Potential ET’ (description of the evapotranspiration computation), and ’Soil Water Dynamics’ (description of the algorithms adopted).

**Table S3.**
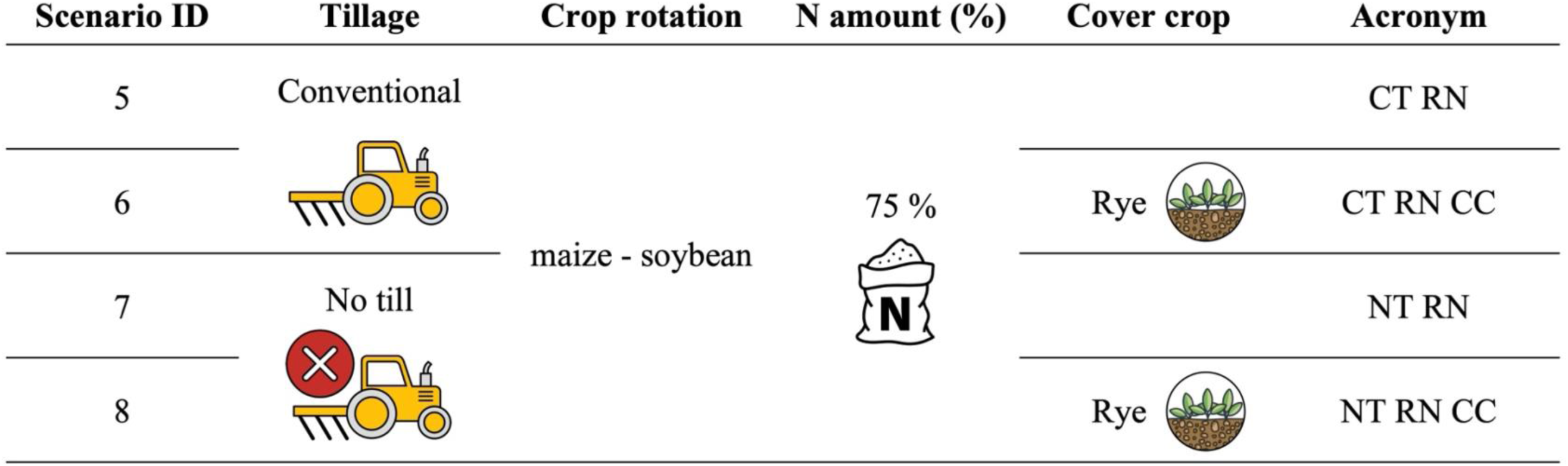
Additional scenarios (5-8) simulated using an ensemble multi-model approach, representing different combinations of management practices involving tillage, nitrogen fertilization reduction and cover crop utilization. The reduced nitrogen fertilization (75%) refers to a reduction of the Midwest state based “business-as-usual” average amount and was applied only for maize. A two-year crop rotation, with maize followed by soybean, was implemented in all scenarios. Scenario acronyms: CT (conventional till), NT (no till), RN (Reduced nitrogen), CC (cover crop).

| Scenario ID | Tillage | Crop rotation | N amount (%) | Cover crop | Acronym |
| --- | --- | --- | --- | --- | --- |
| 5 | Conventional | maize - soybean | 75 % |  | CT RN |
| 6 |  |  |  | Rye | CT RN CC |
| 7 | No till |  |  |  | NT RN |
| 8 |  |  |  | Rye | NT RN CC |

**Table S4.** Crop management practices used to upscale the eight scenarios across the US Midwest. The table provides details for both maize and soybean, including sowing date, plant density, row spacing, sowing depth, harvest dates, and fertilization. Additionally, the same information is reported for the rye cover crop, but only for the scenarios that incorporate it.

| **Crop** | **Sowing date** | **Plant density**<br>**(seeds m<sup>-2</sup>)** | **Row spacing**<br>**(cm)** | **Sowing depth**<br>**(cm)** | **Harvest date** | **Fertilization** |
| --- | --- | --- | --- | --- | --- | --- |
| Maize | Annual NASS planting dates by weather grid | 8 | 76 | 5 | Annual NASS harvest dates by weather grid | Split application with 1/3 at planting as Urea Ammonium Nitrate (UAN) incorporated at 5 cm, 2/3 applied 45 days after planting as UAN incorporated at 2 cm. Total N rate varied by state |
| Soybean | Annual NASS planting dates by weather grid | 40 | 50 | 2.5 | Annual NASS harvest dates by weather grid | No fertilizer added |
| Winter rye<br>(cover crop) | 7 days after harvesting the cash crop | 200 | 15 | 3 | 7 days before planting the cash crop | 30 kg N ha<sup>-1</sup> as Urea Ammonium Nitrate (UAN) at planting incorporated at 2 cm |

**Table S5.** Tillage characteristic used to upscale the eight scenarios across the US Midwest. The table provides details about conventional tillage chronological date order, operation type and depth adopted. No till scenarios do not involve any tillage operations.

| **Tillage** | **Chronological date order** | **Operation type** | **Depth (cm)** |
| --- | --- | --- | --- |
| Conventional | 7 days before cash crop sowing | Chisel plow | 20 |
|  | At cash crop sowing | Field cultivator | 10 |
|  | At cover crop (if present) planting date | Field cultivator | 10 |
| No till |  |  |  |

**Table S6.** Total fertilizer nitrogen rates (kg N ha^-1^) applied per state for the eight scenarios, scaled up across the US Midwest area.

| State | N amount (kg ha<sup>-1</sup>) |
| --- | --- |
| Iowa | 180 |
| Illinois | 204 |
| Indiana | 209 |
| Kansas | 224 |
| Michigan | 170 |
| Minnesota | 183 |
| Missouri | 224 |
| North Dakota | 163 |
| Nebraska | 180 |
| Ohio | 194 |
| South Dakota | 158 |
| Wisconsin | 155 |

**Table S7.** Maize and soybean maturity regions and their corresponding growing degree days (GDD) at maturity used when upscale the eight scenarios across the US Midwest area.

| **Crop** | **Maturity region** | **Cultivar ID** | **GDD to maturity** |
| --- | --- | --- | --- |
| Maize | 1 | 1 | 1146 |
|  | 2 | 2 | 1306 |
|  | 3 | 3 | 1401 |
|  | 4 | 4 | 1466 |
|  | 5 | 5 | 1483 |
|  | 6 | 6 | 1506 |
| Soybean | 1 | 1 | 1295 |
|  | 2 | 2 | 1481 |
|  | 3 | 3 | 1589 |
|  | 4 | 4 | 1702 |
|  | 5 | 5 | 1827 |
|  | 6 | 6 | 1956 |

**Table S8.**
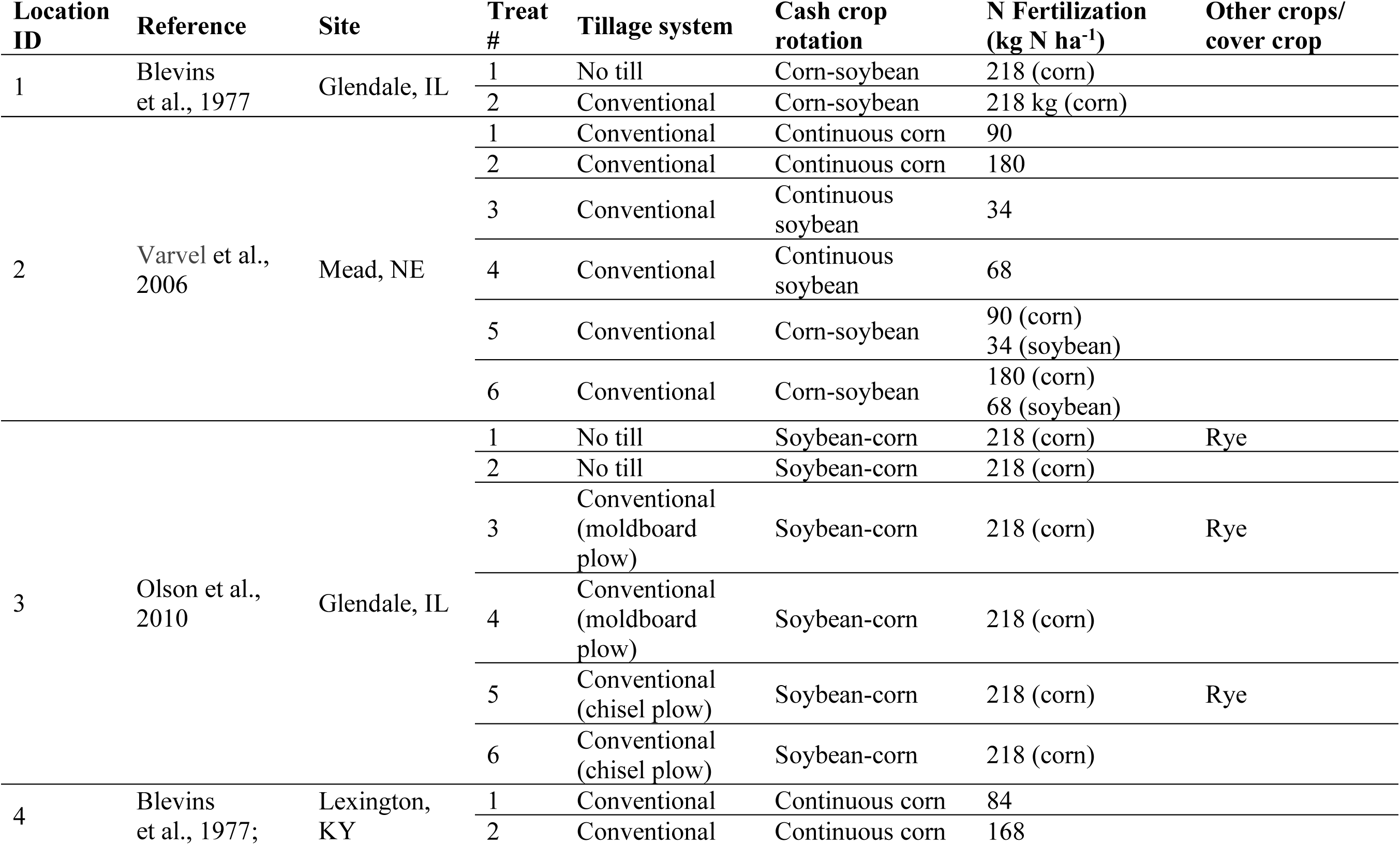
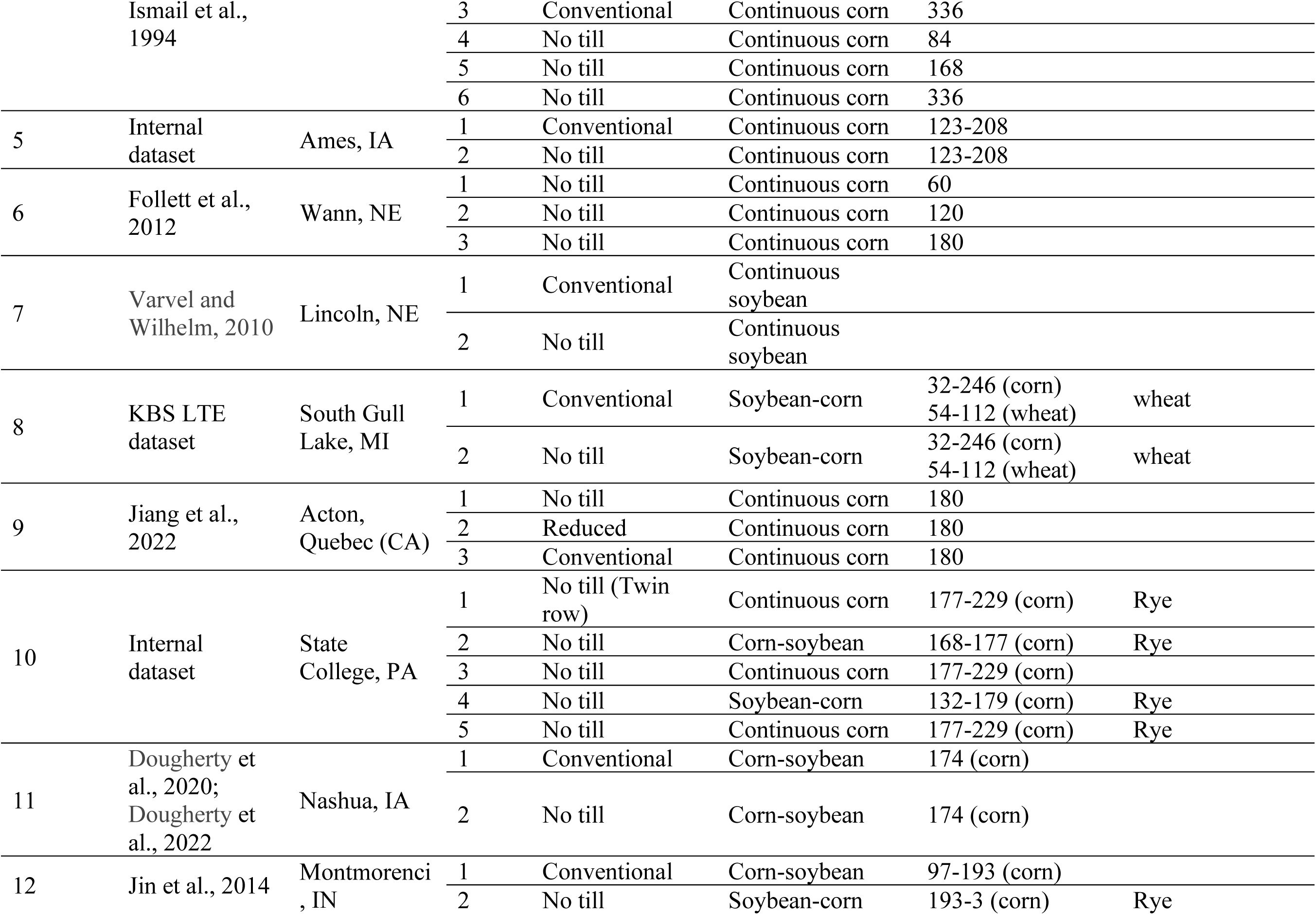
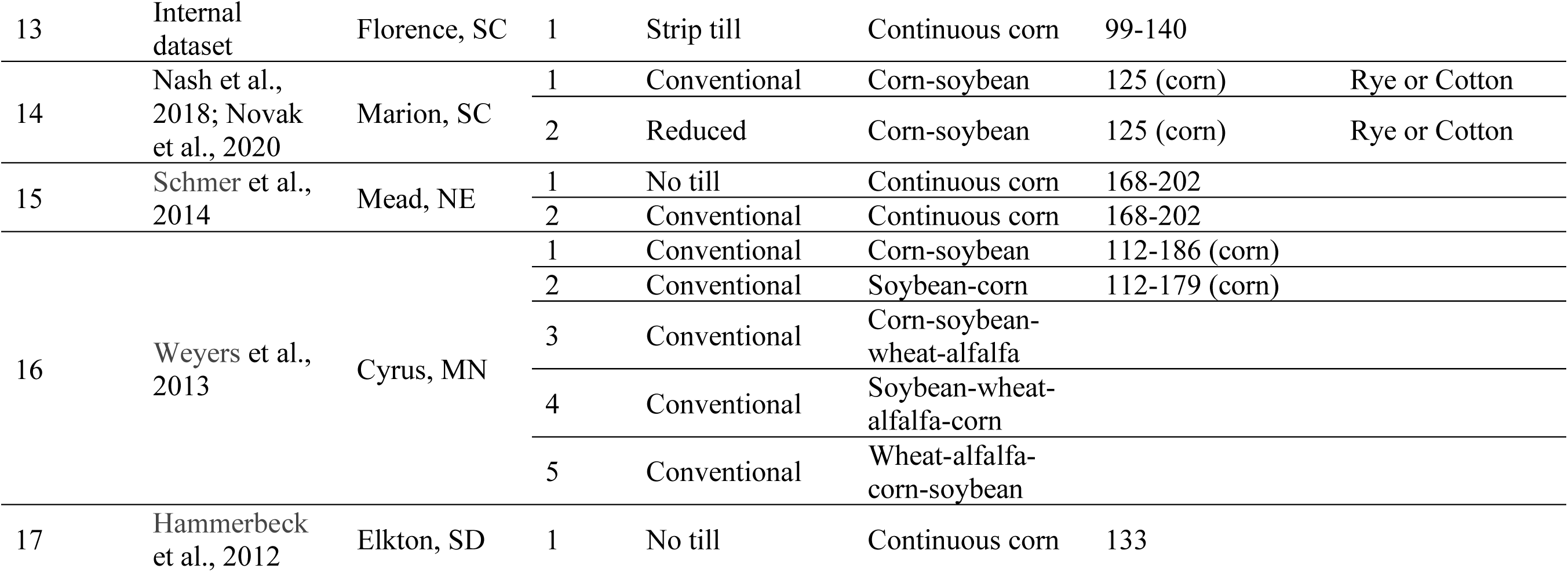
Description of the datasets collected for the Multi-model Ensemble validation. Data are organized by single location and treatments. Table shows the tillage system, main cash crop rotation, and nitrogen fertilization level adopted and presence of additional crops or cover crop.

**Table S9.** Relative Root Mean Square Error (RRMSE, %, equation 3) for yield across 17 locations, three crops (maize, soybean and wheat), and eight models, derived from models’ yield calibration.

| **Location ID** | **Crop** | **Model 1** | **Model 2** | **Model 3** | **Model 4** | **Model 5** | **Model 6** | **Model 7** | **Model 8** |
| --- | --- | --- | --- | --- | --- | --- | --- | --- | --- |
| 1 | Maize | 30 | 28 | 37 | 29 | 26 | 30 | 21 | 27 |
|  | Soybean | 40 | 30 | 25 | 58 | 29 | 38 | 20 | 35 |
| 2 | Maize | 45 | 81 | 48 | 99 | 80 | 46 | 55 | 53 |
|  | Soybean | 17 | 69 | 7 | 61 | 34 | 30 | 33 | 20 |
| 3 | Maize | 26 | 37 | 25 | 41 | 14 | 26 | 22 | 29 |
|  | Soybean | 31 | 33 | 26 | 28 | 28 | 27 | 19 | 23 |
| 4 | Maize | 40 | 33 | 50 | 40 | 28 | 33 | 33 | 31 |
| 5 | Maize | 37 | 28 | 38 | 32 | 30 | 19 | 22 | 12 |
| 6 | Maize | 64 | 43 | 44 | 56 | 64 | 42 | 43 | 41 |
| 7 | Soybean | 24 | 28 | 34 | 51 | 35 | 28 | 31 | 35 |
| 8 | Maize | 36 | 33 | 46 | 25 | 47 | 27 | 31 | 32 |
|  | Soybean | 35 | 29 | 30 | 33 | 48 | 42 | 36 | 36 |
|  | Wheat | 36 | NA | NA | NA | NA | 29 | NA | 27 |
| 9 | Maize | 34 | 39 | 39 | 34 | 39 | 43 | 37 | 37 |
| 10 | Maize | 44 | 34 | 48 | 26 | 37 | 31 | 26 | 31 |
|  | Soybean | 20 | 45 | 46 | 37 | 23 | 37 | 28 | 22 |
| 11 | Maize | 24 | 29 | 35 | 16 | 27 | 14 | 27 | 14 |
|  | Soybean | 38 | 42 | 25 | 25 | 25 | 35 | 28 | 49 |
| 12 | Maize | 36 | 28 | 47 | 29 | 30 | 31 | 21 | 21 |
|  | Soybean | 28 | 31 | 25 | 41 | 38 | 40 | 20 | 17 |
| 13 | Maize | 25 | 21 | 28 | 42 | 38 | 41 | 38 | 19 |
| 14 | Maize | 37 | 33 | 68 | 71 | 55 | 52 | 41 | 51 |
|  | Soybean | 24 | 39 | 32 | 32 | 30 | 32 | 31 | 66 |
| 15 | Maize | 52 | 22 | 33 | 20 | 32 | 28 | 16 | 19 |
| 16 | Maize | 35 | 39 | 31 | 47 | 26 | 24 | 19 | 16 |
|  | Soybean | 35 | 31 | 27 | 30 | 33 | 33 | 20 | 26 |
|  | Wheat | 23 | 85 | 42 | 16 | 26 | 35 | 28 | 30 |
| 17 | Maize | 51 | 45 | 43 | 35 | 37 | 33 | 36 | 47 |
| **Overall** |  | **35** | **38** | **36** | **39** | **35** | **33** | **29** | **31** |

**Table S10.** Full report of the 52 combinations of measured sites (17) and treatments (number depending on the location) under analysis. Table reports the location ID and treatment number (#) as reported in table S8; tillage type (CT = conventional tillage, NT = No till and RT = Reduced tillage); locations coordinates (latitude, longitude and state); duration of the experiment (year); maximum tillage depth (cm) adopted during the experiment; number of crops in the main rotation; cover crop presence; soil organic carbon sampling depth (cm); measured delta SOC changes between samplings (Mg ha^-1^ yr^-1^) computed between every consequent soil sampling; measured weighed mean delta SOC change (Mg ha^-1^ yr^-1^); MME weighted mean delta SOC change (Mg ha^-1^ yr^-1^) and consistency of the measured weighed mean delta SOC change with literature ranges. Single combinations are grouped by literature consistency, tillage and cover crop presence.

| Location ID | Treat # | Tillage | Coordinates | Duration (years) | Max tillage depth (cm) | N* of crops in rotation | Cover crop | SOC sampling depth (cm) | Measured delta SOC changes between consequent samplings ($\text{Mg ha}^{-1} \text{ yr}^{-1}$) | Measured weighted mean delta SOC change ($\text{Mg ha}^{-1} \text{ yr}^{-1}$) | MME weighted mean delta SOC change ($\text{Mg ha}^{-1} \text{ yr}^{-1}$) | Literature values consistency |
| --- | --- | --- | --- | --- | --- | --- | --- | --- | --- | --- | --- | --- |
| 1 | 2 | CT | 37.432, -88.667 (IL) | 24 | 15 | 3 | none | 15 | -0.7, -0.083 | -0.39 | 0.15 | YES |
| 2 | 1 | CT | 41.162, -96.416 (NE) | 18 | 20 | 1 | none | 30 | -0.155, -0.378, -0.375 | -0.28 | 0.05 | YES |
| 2 | 2 | CT | 41.162, -96.416 (NE) | 18 | 20 | 1 | none | 30 | 0.035, -0.27, -0.398 | -0.16 | 0.05 | YES |
| 2 | 3 | CT | 41.162, -96.416 (NE) | 18 | 20 | 1 | none | 30 | -0.446, -0.203, -0.323 | -0.34 | -0.14 | YES |
| 2 | 4 | CT | 41.162, -96.416 (NE) | 18 | 20 | 1 | none | 30 | -0.519, -0.152, -0.03 | -0.29 | -0.13 | YES |
| 2 | 5 | CT | 41.162, -96.416 (NE) | 18 | 20 | 2 | none | 30 | -0.274, -0.117, -0.042 | -0.17 | -0.10 | YES |
| 2 | 6 | CT | 41.162, -96.416 (NE) | 18 | 20 | 2 | none | 30 | -0.378, -0.007, 0.037 | -0.16 | -0.09 | YES |
| 3 | 6 | CT | 37.434, -88.667 (IL) | 11 | 15 | 2 | none | 15 | 2.602, -1.75, -0.2, 0.867 | 0.04 | 0.32 | YES |
| 4 | 1 | CT | 38.129, -84.497 (KY) | 20 | 20 | 2 | rye | 30 | -2.423, 1.618 | 0.27 | 0.34 | YES |
| 4 | 2 | CT | 38.129, -84.497 (KY) | 20 | 20 | 2 | rye | 30 | -2.085, 1.459 | 0.27 | 0.50 | YES |
| 3 | 5 | CT | 37.434, -88.667 (IL) | 11 | 15 | 3 | rye | 15 | 2.702, -1.35, 0.3, 1.067 | 0.35 | 0.56 | YES |
| 5 | 2 | NT | 42.018, -93.764 (IA) | 5 | 10 | 1 | none | 15 | -0.3, -1.101, 1.501 | -0.10 | 0.21 | YES |
| 1 | 1 | NT | 37.431, -88.665 (IL) | 24 | 5 | 3 | none | 15 | -0.142, -0.042 | -0.09 | 0.24 | YES |
| 6 | 1 | NT | 41.168, -96.411 (NE) | 14 | NA | 1 | none | 30 | 2.115, -0.08, 0.705 | 0.89 | 0.37 | YES |
| 6 | 2 | NT | 41.168, -96.411 (NE) | 14 | NA | 1 | none | 30 | 2.772, 2.446, -1.046 | 1.15 | 0.57 | YES |
| 6 | 3 | NT | 41.168, -96.411<br>(NE) | 14 | NA | 1 | none | 30 | 1.875, 1.79, -0.605 | 0.86 | 0.59 | YES |
| 3 | 2 | NT | 37.434, -88.667 (IL) | 11 | NA | 2 | none | 15 | 1.451, -1.2, 0.15,<br>2.168 | 0.45 | 0.31 | YES |
| 7 | 2 | NT | 40.848, -96.467<br>(NE) | 5 | 4 | 1 | none | 30 | 0.08 | 0.08 | 0.06 | YES |
| 8 | 2 | NT | 42.395, -85.374<br>(MI) | 11 | 5 | 3 | none | 25 | 1.082, -2.041, 1.502,<br>0,<br>-1.044, 2.316, -1.708,<br>1.584, 3.845, -5.537,<br>-0.302, 7.605 | 0.72 | 0.51 | YES |
| 9 | 1 | NT | 45.5, -72.416 (QC<br>CA) | 22 | 5 | 1 | none | 20 | -0.038, 2.841, -1.191,<br>-3.328, 15.483,<br>13.173, 0.65 | 0.74 | 0.20 | YES |
| 10 | 4 | NT | 40.791, -77.858<br>(PA) | 4 | NA | 3 | rye | 30 | -0.12 | -0.12 | 0.97 | YES |
| 3 | 1 | NT | 37.434, -88.667 (IL) | 11 | NA | 3 | rye | 15 | 1.701, -0.925,<br>-0.999, 0.6 | -0.05 | 0.76 | YES |
| 11 | 2 | NT | 42.936, -92.569 (IA) | 10 | 5 | 3 | rye | 15 | 0.49 | 0.49 | 0.28 | YES |
| 10 | 1 | NT | 40.791, -77.858<br>(PA) | 4 | NA | 2 | rye | 30 | 0.56 | 0.56 | 1.31 | YES |
| 10 | 2 | NT | 40.791, -77.858<br>(PA) | 4 | NA | 3 | rye | 30 | 0.40 | 0.40 | 1.12 | YES |
| 10 | 5 | NT | 40.791, -77.858<br>(PA) | 4 | NA | 2 | rye | 30 | 0.43 | 0.43 | 1.25 | YES |
| 4 | 4 | NT | 38.129, -84.497<br>(KY) | 20 | NA | 2 | rye | 30 | -1.345, 1.284 | 0.40 | 0.75 | YES |
| 4 | 5 | NT | 38.129, -84.497<br>(KY) | 20 | NA | 2 | rye | 30 | -1.267, 1.30 | 0.44 | 0.87 | YES |
| 4 | 6 | NT | 38.129, -84.497<br>(KY) | 20 | NA | 2 | rye | 30 | 0.247, 1.278 | 0.93 | 0.93 | YES |
| 12 | 2 | NT | 40.472, -86.998 (IN) | 7 | 5 | 3 | rye | 10 | 1.156, -1.801,<br>3.102, -1.697 | 0.36 | 0.35 | YES |
| 13 | 1 | RT | 34.283, -79.741<br>(SC) | 4 | 30 | 1 | none | 15 | -0.582, -0.368,<br>-0.17, 0.036 | -0.27 | 0.05 | YES |
| 14 | 2 | RT | 34.171, -79.441<br>(SC) | 11 | 40 | 4 | rye | 15 | 1.866, -2.402, 3.502,<br>-0.4, -0.499, 0.1,<br>2.402,<br>-3.602 | 0.47 | 0.21 | YES |
| 11 | 1 | CT | 42.936, -92.569 (IA) | 10 | 20 | 2 | none | 15 | 0.75 | 0.75 | 0.29 | NO |
| 15 | 2 | CT | 41.162, -96.412<br>(NE) | 13 | 15 | 1 | none | 30 | -0.405, -2.022 | -0.88 | 0.17 | NO |
| 5 | 1 | CT | 42.017, -93.764 (IA) | 5 | 20 | 1 | none | 15 | -0.933, -0.901, -1.001 | -0.94 | 0.15 | NO |
| 12 | 1 | CT | 40.472, -86.998 (IN) | 7 | 25 | 2 | none | 10 | -0.475, -0.998 | -0.58 | 0.14 | NO |
| 16 | 1 | CT | 45.7, -95.8 (MN) | 5 | 23 | 2 | none | 15 | 2.58 | 2.58 | -0.01 | NO |
| 16 | 2 | CT | 45.7, -95.8 (MN) | 5 | 23 | 2 | none | 15 | 1.20 | 1.20 | -0.03 | NO |
| 7 | 1 | CT | 40.848, -96.467 (NE) | 5 | 20 | 1 | none | 30 | -0.55 | -0.55 | -0.11 | NO |
| 16 | 3 | CT | 45.7, -95.8 (MN) | 5 | 23 | 4 | none |  | 1.82 | 1.82 | 0.00 | NO |
| 8 | 1 | CT | 42.395, -85.374 (MI) | 31 | 25 | 3 | none | 25 | 0.522, -1.938, 2.214, -1.378, -3.392, 1.782, 0.569, -2.079, 8.458, -6.709, -1.509, 6.504 | 0.56 | 0.45 | NO |
| 16 | 4 | CT | 45.7, -95.8 (MN) | 5 | 23 | 4 | none | 15 | 1.91 | 1.91 | 0.06 | NO |
| 9 | 3 | CT | 45.5, -72.416 (QC CA) | 22 | 20 | 1 | none | 20 | -1.814, 6.27, -13.207, 0.281, -7.344, 0.432 | -0.61 | -0.38 | NO |
| 16 | 5 | CT | 45.7, -95.8 (MN) | 5 | 18 | 4 | none | 15 | 2.49 | 2.49 | 0.19 | NO |
| 3 | 4 | CT | 37.434, -88.667 (IL) | 11 | 15 | 2 | none | 15 | -3.102, -0.4, 0.05, 0.534 | -0.56 | 0.28 | NO |
| 14 | 1 | CT | 34.171, -79.441 (SC) | 11 | 40 | 4 | rye | 15 | 0.3, 2.794, -1.501, 2.902, -2.202, 1.996, 2.702, -1.501, -0.801 | 0.50 | 0.31 | NO |
| 3 | 3 | CT | 37.434, -88.667 (IL) | 11 | 15 | 3 | rye | 15 | -2.001, -0.725, -0.55, 0.634 | -0.56 | 0.46 | NO |
| 4 | 3 | CT | 38.129, -84.497 (KY) | 20 | 20 | 2 | rye | 30 | -1.697, | 0.60 | 0.55 | NO |
| 15 | 1 | NT | 41.162, -96.412 (NE) | 13 | 5 | 1 | none | 30 | -0.183, -1.43 | -0.55 | 0.24 | NO |
| 10 | 3 | NT | 40.791, -77.858 (PA) | 4 | NA | 1 | none | 30 | -0.96 | -0.96 | 0.98 | NO |
| 17 | 1 | NT | 44.19, -96.46 (SD) | 5 | NA | 1 | none | 30 | 2.77, 1.828, 12.14, -0.773 | 3.99 | 0.69 | NO |
| 9 | 2 | RT | 45.5, -72.416 (QC CA) | 22 | 15 | 1 | none | 20 | -0.497, 3.946, -14.57, -0.841, 18.562, 8.139, 2.186 | 1.14 | 0.09 | NO |

## Dataset S1 (separate file)

Dataset_S1.xlsx contains eight detailed tables, one for each CSM, describing the calibrated parameters used in each of the 17 experimental sites during the MME testing phase. These parameters were used to set each the yield of each crop cultivar involved in the rotation across the sites. to The columns in the tables include parameter names, descriptions, units of measurement, calibration methods, uncalibrated default values, and calibrated values for each location. Each row corresponds to a specific parameter, illustrating how its value changes from the uncalibrated to the calibrated phase across different sites.

## Notes

### Competing Interest Statement

Bruno Basso is a cofounder of CIBO Technologies. Keith Paustian and Yao Zhang have financial interest in Indigo Ag. The other coauthors declare no conflict of interest.

